# The G1/S transition is promoted by Rb degradation via the E3 ligase UBR5

**DOI:** 10.1101/2023.10.03.560768

**Authors:** Shuyuan Zhang, Lucas Fuentes Valenzuela, Evgeny Zatulovskiy, Lise Mangiante, Christina Curtis, Jan M. Skotheim

## Abstract

Mammalian cells make the decision to divide at the G1/S transition in response to diverse signals impinging on the retinoblastoma protein Rb, a cell cycle inhibitor and tumor suppressor. Rb is inhibited by two parallel pathways. In the canonical pathway, Cyclin D-Cdk4/6 kinase complexes phosphorylate and inactivate Rb. In the second, recently discovered pathway, Rb’s concentration decreases during G1 to promote cells progressing through the G1/S transition. However, the mechanisms underlying this second pathway are unknown. Here, we found that Rb’s concentration drop in G1 and recovery in S/G2 is controlled by phosphorylation-dependent protein degradation. In early G1 phase, un- and hypo-phosphorylated Rb is targeted by the E3 ligase UBR5. *UBR5* knockout cells have higher Rb concentrations in early G1, exhibit a lower G1/S transition rate, and are more sensitive to Cdk4/6 inhibition. This last observation suggests that UBR5 inhibition can strengthen the efficacy of Cdk4/6 inhibitor-based cancer therapies.

## Introduction

The decision to divide often takes place in the G1 phase of the cell cycle and occurs in response to diverse input signals. Once taken, the decision to initiate DNA replication and divide is difficult to reverse despite changes to the input signals ^1,2^. From a molecular point of view, the commitment point at the G1/S transition in response to growth signals corresponds to the hyper- phosphorylation and inactivation of the transcriptional inhibitor Rb, the retinoblastoma protein ^1,3,4^. Hyper-phosphorylation of Rb frees the activating E2F transcription factors to drive expression of the Cyclin E and A, which can form complexes with the cyclin-dependent kinase Cdk2. Active Cyclin E/A-Cdk2 complexes then maintain Rb hyper-phosphorylation so that E2F- dependent transcription remains active throughout S phase ^5^. While the molecular basis of the commitment point to cell division is increasingly well understood^1,2,6–8^, we know much less about how the upstream input signals transmit quantitative information to the decision point.

Two distinct input pathways regulating the G1/S transition operate by inactivating Rb. The first pathway, which is the canonical pathway, acts through Cyclin-Cdk complexes phosphorylating Rb. Decades of work has gradually unveiled the molecular mechanisms underlying this pathway^5,9–12^. The upstream growth factors initiate signals that increase the expression of Cyclin D ^13^, which primarily forms a complex with the cyclin-dependent kinases Cdk4 and Cdk6 ^14^. Cyclin D- Cdk4/6 complexes then initiate the phosphorylation of Rb, possibly through hypo- or mono- phosphorylation ^15,16^. The hypo-phosphorylation of Rb likely shifts the dissociation constant (Kd) of Rb with E2F, thus promoting the G1/S transition. Once Rb is hyper-phosphorylated, possibly by the increasing Cyclin E-Cdk2 activity and the initial Cyclin D-Cdk4/6 activity, it is fully inactivated so that E2F can drive the E2F-dependent S phase transcription program. This Rb phosphorylation pathway is frequently hijacked in cancers to drive cell proliferation ^17–21^. For example, Cyclin D amplification is a frequent mutation in breast cancer patients and is associated with shorter relapse free survival ^17,22^. Moreover, increasing the Cyclin D-Cdk4 activity by transducing cells with a *CDK4* construct is a common approach to immortalize cells *in vitro* ^23,24^. In fact, for many cancer cell lines and immortalized cell lines, Rb is immediately hyper- phosphorylated after cell birth due to a high Cdk activity likely arising both from mutations selected for proliferation and from the rich *in vitro* culture environment ^2,25^.

A second pathway that inactivates Rb, which we recently identified, operates through decreasing the concentration of the Rb protein during cell growth in G1 ^26–28^. Specifically, the total amount of Rb protein stays relatively constant throughout G1, while the cell is growing bigger, so that its concentration decreases. In contrast, the concentrations of G1/S activators, including Cyclin D and E2F, stay constant during early to mid G1. These differential effects on protein concentration in G1 phase drive relative changes in the activities of the cell cycle inhibitor (Rb) and activators (E2F, Cyclin D) to favor progression through the G1/S transition.

Thus, our current model is that two Rb-inactivation pathways cooperate to activate E2F- dependent transcription and initiate the cell cycle^26–28^ (Figure 1A). Namely, Cyclin D-dependent Rb hypo-phosphorylation shifts the dissociation constant (Kd) of Rb with E2F so that the decreasing Rb concentration can drop below Kd to release active E2F. Following the G1/S transition, Rb’s concentration increases during S/G2/M to reset for the next cell cycle. Unlike the Cdk-dependent pathway, the Rb-dilution pathway does not depend on any increased Cdk activity and can serve as a parallel mechanism to inactivate Rb and promote the G1/S transition. Although several molecular mechanisms underlying Cyclin D synthesis and Rb phosphorylation have been elucidated ^15,29^, the molecular mechanism underlying the second pathway that decreases Rb concentration through G1 is unknown.

**Figure 1.**
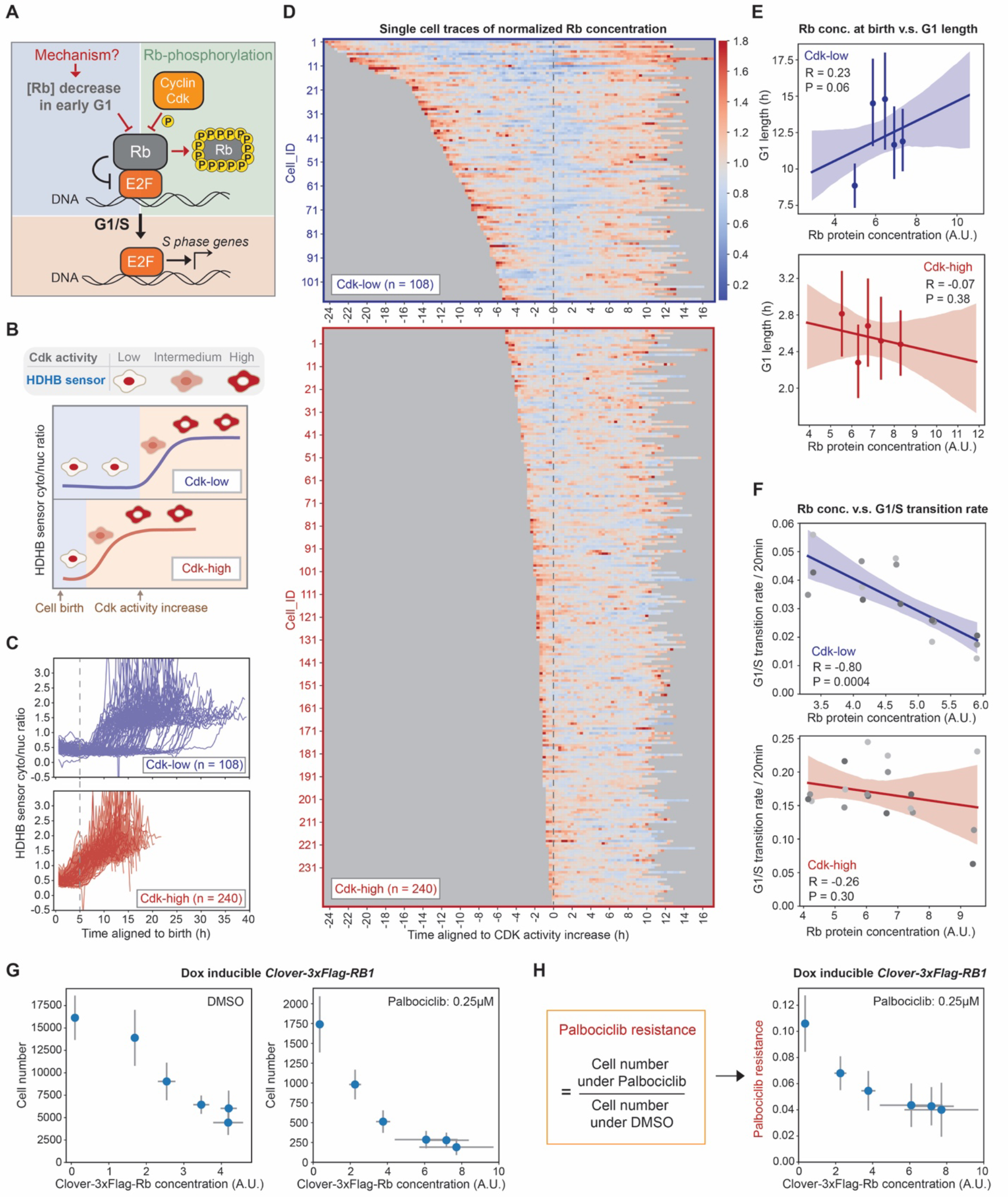
Rb concentration dynamics regulate cell cycle progression in Cdk-low cells A. Schematic illustration of two parallel pathway acting through Rb concentration and Rb phosphorylation to promote the G1/S transition. **B**. Schematic illustration of the Cdk activities in Cdk-low and Cdk-high categories of cells. Cdk activity is measured using the HDHB sensor whose translocation to the cytoplasm from the nucleus is driven by Cdk activity. Cdk activation is defined using the inflection point in the cytoplasm-to-nuclear ratio. **C.** Fluorescent traces showing the HDHB cytoplasm-to-nucleus intensity ratio from HMEC-hTERT1 cells expressing endogenously tagged *RB1-3xFLAG-Clover-sfGFP* and the HDHB Cdk sensor ^25^. Cdk-high cells are defined as those activating Cdk within 5 hours of their birth, while Cdk-low cells activate Cdk later. **D.** Individual fluorescent traces from the cells in (C). Upper panel shows Cdk-low cells, and the lower panel shows Cdk-high cells. Traces are aligned by the initial translocation of the HDHB Cdk sensor. The trace color represents the normalized Rb protein concentration (normalized to the mean Rb concentration of the cells in the same group). **E**. The correlations between Rb concentration at birth and G1 length in Cdk-low and Cdk-high cells. **F**. The correlations between Rb concentration and G1/S transition rate in Cdk-low and Cdk-high cells. **G.** Cell number of HMEC cells treated with 0.25 µM Palbociclib for 72 hours. Cells were plated in different doses of doxycycline to induce exogenous Clover-3xFlag-Rb. Drug treatment started the next day and lasted for 72 hours. Then, cells were fixed and the cell number in each well was measured. **H**. Normalized cell number of the cells treated as described in panel (**G**). Normalized cell number is the cell number following Palbociclib treatment divided by the cell number under DMSO control treatment. n = 3 biological replicates, the error bars indicate standard deviation.

Here, we sought to determine the molecular mechanisms underlying Rb’s concentration decrease through G1 (Figure 1A), an important signal triggering the G1/S transition. We found that Rb is consistently synthesized throughout the cell cycle, but its degradation is cell cycle dependent. Specifically, Rb is targeted for degradation in G1 by the E3 ligase UBR5 and stabilized by hyper-phosphorylation at the G1/S transition. A mathematical model shows that this mechanism is sufficient to explain the observed concentration dynamics through the cell cycle. Disruption of this Rb degradation mechanism via UBR5 deletion decreases the G1/S transition rate and sensitizes cells to chemical inhibition of Cdk4/6 activity. This last observation provides implications for improving the efficacy of Cdk4/6 inhibitor based therapies through targeting UBR5. Indeed, we observe frequent amplification of *UBR5* in breast cancer patients that is associated with worse prognosis.

## Results

### Rb concentration dynamics regulate cell cycle progression in cells born with low Cdk activity

While the decreasing Rb concentration in G1 drove cell cycle progression in some cells ^26,27^, many cell lines appeared to not respond to changes in Rb dosage in terms of their proliferation rate ^16,30^. We therefore sought to further test the effect of Rb concentration on cell cycle progression and identify the reason behind this apparent discrepancy. To do this, we performed live-cell imaging on *HMEC-hTERT1* (telomerase-immortalized human mammary epithelial, abbreviated as HMEC) cells expressing endogenously tagged *RB1-3xFLAG-Clover-sfGFP* and the HDHB Cdk activity sensor ^25^. The nuclear-to-cytoplasmic translocation of this fluorescent sensor marks the transition point in mid/late-G1 when Cdk activity abruptly increases (Figure 1B) ^1,25^. Based on how quickly the Cdk activity rises after birth, we classified the cells into two categories: Cdk-high cells and Cdk-low cells based on whether or not Cdk activity had risen 5 hours after birth (Figure 1B, C; Figure S1C). This classification has been used in other studies that applied the same Cdk sensor on MCF10A cells ^25^. When we aligned the cell traces to the inflection point of the HDHB sensor that marks Cdk activity increase, we found that the concentration of Rb continuously decreases during early- to mid-G1 phase in the Cdk-low cells, and then increases through the remainder of the cell cycle (Figure 1D; Figure S1A, B, D-G).

However, Cdk-high cells do not exhibit decreasing Rb concentration dynamics in G1 (Figure 1D; Figure S1A, B, D-G). Thus, an Rb concentration decrease in G1 could only regulate the cell cycle in Cdk-low cells, but not in Cdk-high cells, where the concentration decrease does not take place and Rb is likely rapidly inactivated.

To test if the decrease in Rb concentration promotes cell cycle entry in Cdk-low cells, we examined the relationship between Rb concentration and G1 length and G1/S transition rate. We found that the Rb concentration at birth is positively correlated with G1 length (Figure 1E), and the Rb concentration is anti-correlated with the G1/S transition rate (Figure 1F). That these correlations are significant in Cdk-low cells but not in Cdk-high cells is consistent with Rb concentration regulating the G1/S transition only when Cdk activity is low. To further test this by exogenously controlling the concentration of Rb, we used HMEC cells containing a Dox-inducible allele of *RB1*. Increasing Rb concentration not only suppresses cell proliferation, but also further sensitizes cells to treatment by Cdk4/6 inhibitors (Figure 1G,H; Figure S2). Taken together, these experiments are consistent with our previous work and a recent study reporting that decreasing Rb concentration in G1 plays a crucial role in driving cells into the cell cycle in the absence of Cdk4/6 activity and facilitates the adaptation to chemical Cdk4/6 inhibitors ^31^.

Our findings demonstrate that Rb concentration more significantly impacts the cell cycle of cells with initially low Cdk activity. This makes sense because cells born with high Cdk activity likely quickly inactivate Rb via phosphorylation. This observation also explains why in many experiments overexpression of Rb does not have a big impact on cell proliferation ^16,32^. This is likely because the majority of cells *in vitro* cultured cell lines, especially cancer cell lines, are in the Cdk-high category ^25,32^. Even in the non-cancer cell lines, like the telomerase immortalized HMEC cells we used here, the majority of cells are born with high Cdk activity. However, this is not true in the cells growing *in vivo*. For example, the epidermal stem cells in mouse skin tissue have a much longer G1 duration than *in vitro* cultured cell lines and are not born with high Cdk activity^28,33^. Therefore, the Rb concentration is likely to have a bigger impact on cell cycle progression under *in vivo* settings.

### Rb concentration dynamics are driven by cell cycle-dependent protein degradation

To identify the mechanism regulating Rb concentration dynamics through the cell cycle, we examined both the synthesis and degradation of Rb protein in different cell cycle phases. We measured the mRNA concentration of *RB1* in different cell cycle phases by using flow cytometry to sort HMEC cells expressing FUCCI ^34^ cell cycle reporters into G1 and S/G2 populations. We then performed qPCR and mRNA-Seq to measure *RB1* mRNA expression. The results showed that the *RB1* mRNA concentration did not significantly increase in S/G2 phase (Figure 2A, Figure S3A). We found a similar result when calculating *RB1* mRNA concentrations in different cell cycle phases using a published MERFISH dataset ^35^ (Figure S3B). This indicates that Rb concentration dynamics are controlled by post-transcriptional mechanisms. To further investigate the synthesis dynamics of Rb protein, we measured the translation efficiency of *RB1* mRNA by performing a RIP (RNA-binding protein immunoprecipitation) assay against the translation initiation factor eIF4E^77^. The relative translation efficiency is calculated by dividing the bound fraction of *RB1* with the bound fraction of housekeeping genes (*Actin* and *GAPDH*). Using this method, we found the relative translation efficiency of *RB1* was similar in sorted G1 and S/G2 cells as well as in asynchronously dividing and G1 arrested cells (Palbociclib treatment) (Figure 2B; Figure S3C). Taken together, our data indicate that Rb synthesis is not primarily responsible for its cell cycle dynamics.

**Figure 2.**
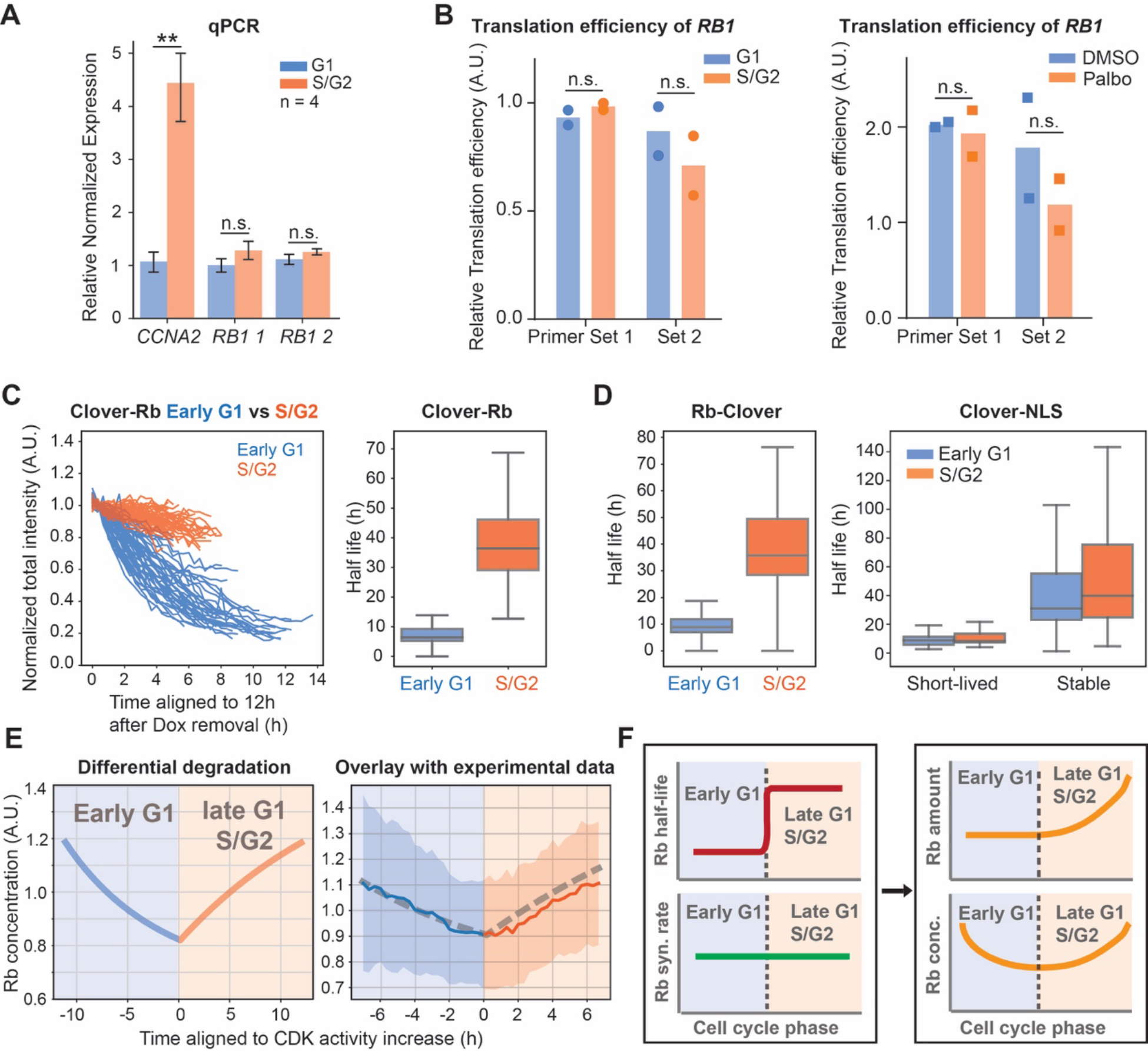
Rb concentration dynamics are driven by cell cycle-dependent protein degradation. **A.** qPCR (n = 4) measurements of the *RB1* mRNA concentration in G1 and S/G2 HMEC cells sorted using a FUCCI cell cycle reporter. **B**. Translation efficiency (TE) of *RB1* was determined in HMEC cells in G1 and S/G2 that had been sorted using a FUCCI marker (left panel), and in HMEC cells treated with DMSO or Palbociclib (1µM) (right panel). TE was measured using a RIP assay (RNA binding protein immunoprecipitation) by pulling down eIF4E. TE is calculated by dividing the eIF4E bound fraction of *RB1* mRNA to the eIF4E bound fraction of *GAPDH* and *Actin* mRNAs. Bars denote mean values and dots denote each replicate experiment. **C.** (Left panel) The degradation traces of Clover-3xFlag-Rb protein after Dox withdrawal. The traces were classified into early G1 phase or S/G2 phase based on a FUCCI cell cycle marker and cell cycle phase duration. (Right panel) Distribution of half-lives estimated from exponential fits. Box plot indicates 5th, 25th, median, 75th and 95th percentiles. **D.** Same half-life measurement as in panel (**C**), but using a C-terminally tagged Rb-3xFlag-Clover (Left panel), a short-lived Clover- NLS, and a stable Clover-NLS (Right panel). **E**. Left panel: Rb concentration dynamics assuming that its degradation rate decreases by 80% at the G1/S transition as measured by live imaging (see Figure 2C) and its synthesis rate does not change (left panel). Right panel shows the overlay of the model based on regulated Rb degradation with the experimental data from Figure S1D (Cdk-low cells). Rb concentration is normalized to the mean. **F.** Model schematic: differential degradation of Rb in G1 and S/G2 phases of the cell cycle drive its concentration dynamics.

Having found that Rb’s cell cycle dynamics were not primarily due to transcription or translation mechanisms, we next sought to test if protein degradation was responsible. To do this, we utilized a doxycyclin (Dox)-inducible system in which cells conditionally express Clover-3xFlag- tagged Rb (*TRE-Clover-3xFlag-Rb*) or Clover-NLS (*TRE-Clover-NLS*) upon Dox treatment (1µg/mL). After 36h of Dox treatment, we withdrew Dox and monitored the decrease of the Clover fluorescence signal using live cell imaging (Figure S4A). Since the cells also express a FUCCI cell cycle marker, we can separately assess protein degradation taking place in G1 and S/G2 phases of the cell cycle (Figure S4A). By fitting the degradation traces using a simple exponential decay function, we obtained the half-life of Clover-3xFlag-Rb protein in different cell cycle phases for each cell. Rb half-life in early G1 (median 6.4h; 75% range 5.3h-9.4h) is significantly shorter than it is in S/G2 (median 37.3h, 75% range 29.2h-45.4h) (Figure 2C). The Rb tag location does not affect its half-life since a C-terminally tagged Rb protein (Rb-3xFlag- Clover) behaved similarly to the N-terminally tagged version (Figure 2D). The changing protein stability in G1 compared to S/G2 phases was specific to Rb as the short-lived Clover-NLS protein and stable Clover-NLS protein expressed with the same Dox-inducible system both had half-lives that did not change through the cell cycle (Figure 2D; Figure S4B, C). Thus, these results suggest a model where the Rb protein’s cell cycle dynamics are due to its degradation in G1 and stabilization at the G1/S transition.

Having established that the regulation of Rb stability is most likely responsible for its cell cycle dynamics, we sought to test if this differential degradation of Rb is sufficient to give rise to the observed dynamics. To do this, we generated a mathematical model where only the half-life of the Rb protein changed through the cell cycle, while the synthesis rate remained constant (see methods). This simple model revealed that regulated degradation was sufficient to generate the cell-cycle dependent Rb concentration dynamics we observed, while the modest upregulation in the synthesis rate was insufficient (Figure 2E; Figure S4D). Note that the Rb protein ‘dilution phenomenon’ that we previously observed in ^26^, in which the total Rb protein amount is kept at a constant level during early G1 while the cell size is growing bigger, is a result of the changing balance between Rb synthesis and Rb being more actively degraded during early G1. From the model, we also examined the dynamics of total Rb protein amount, and we found that, with the experimental parameters for synthesis and degradation rates, the Rb protein amount is relatively constant in early G1 phase (Figure S4E). This shows that the observed dilution in G1 can be explained by Rb’s degradation rate being similar to its synthesis rate in G1 while cell size increases. Taken together, our data and analysis indicate that the cell cycle-dependent regulation of Rb stability is primarily responsible for its cell cycle dynamics (Figure 2F).

### Rb is stabilized via phosphorylation by Cdk

Having found that Rb is stabilized at the G1/S transition, we next sought to identify the molecular mechanism. One of the most prominent molecular changes occurring at the G1/S transition is the phosphorylation of Rb by Cyclin-Cdk complexes. We therefore sought to examine how Rb phosphorylation affected its half-life. To do this, we first stained asynchronous HMEC cells with phospho-Rb (S807/811) and total Rb antibodies. We then calculated the Rb concentration in the low phospho-Rb G1 population (low pRb G1), the high phospho-Rb G1 population (high pRb G1), and the S/G2 population. The Rb concentration is lower in early G1 when it is not hyper-phosphorylated and then begins to recover in late G1, when Rb is phosphorylated (Figure 3A). Similar results were obtained when G1 was partitioned into early and late phases using a recently published live-cell Cdk activity sensor (KTR sensor) based on the C-terminal part of Rb (886-928aa of Rb) ^36^ (Figure S5A). These immunofluorescence data support the model where the Rb concentration decrease in G1 phase is reversed upon its phosphorylation. Consistently, when cells are arrested in G1 by treating them with the Cdk4/6 inhibitor Palbociclib (1µM) for 24 hours, the Rb protein concentration drops by ∼75% (Figure 3B) even though the mRNA concentration is only reduced by ∼15% (Figure S5B). This is consistent with published results showing significant Rb protein drops when cells are exposed to Cdk4/6 inhibitors^37^. Furthermore, in cells expressing a Dox-inducible Clover-3xFlag-Rb protein, Palbociclib treatment led to a significant decrease in the concentration of this ectopically expressed protein but not the corresponding mRNA (Figure S5C). Taken together, these data suggest that the phosphorylation of Rb by Cdk mediates its stabilization. To test this model, we used the HDHB Cdk sensor to categorize cells into Cdk-low and Cdk-high populations before measuring the half-lives of Clover-Rb and Clover-NLS (Figure S5D, E). As anticipated, we found that Rb was degraded much more rapidly in the Cdk-low population.

**Figure 3.**
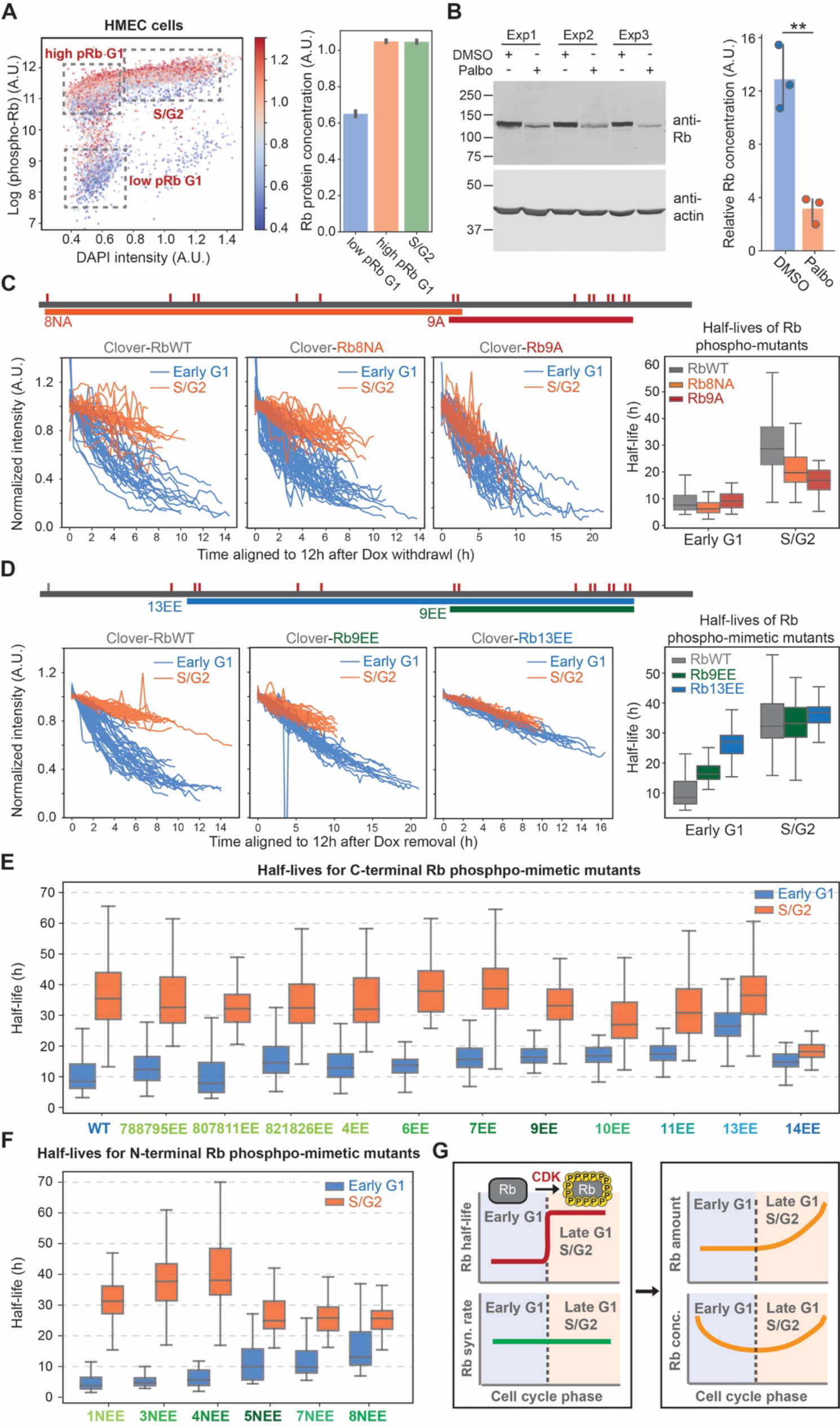
Rb protein is stabilized via phosphorylation by Cyclin-Cdk A. Rb concentration in different phospho-Rb populations. HMEC cells were stained with phospho-Rb (S807/811) and Rb antibodies. (Left panel) phospho-Rb intensity is plotted against DNA content (DAPI intensity). The color indicates Rb concentration. (Right panel) quantification of Rb concentrations in different phospho-Rb populations gated using the indicated boxes in the left panel. Bar plots indicate the mean and 95% confidence interval. **B.** Immunoblot of Rb after DMSO or Palbociclib (1µM) treatment for 24 hours. The quantification of relative Rb concentration (normalized to actin intensity) is shown on the right. **C.** Top panel shows the schematic of Rb phospho-site mutants. Small red lines indicate the location of Cdk phosphorylation sites. Lower panel shows the degradation traces for Clover-3xFlag-RbWT, Clover-3xFlag-Rb8NA, and Clover-3xFlag-Rb9A as well as the corresponding distributions of half-lives. **D.** Top panel shows the schematic of the Rb phospho-mimetic mutants. Lower panel shows the degradation traces for Clover-3xFlag-RbWT, Clover-3xFlag-Rb9EE, and Clover- 3xFlag-Rb13EE, as well as the corresponding distributions of half-lives. **E-F.** Half-life distributions for all the Rb phospho-mimetic mutants. **G.** Model schematic: Rb is stabilized in late G1 and S/G2 phases by Cdk phosphorylation.

To further investigate how Rb phosphorylation on different Cdk phosphorylation sites affects its half-life, we used the Dox-inducible system to express a series of Rb variants in which the Cdk phosphorylation sites were either substituted with non-phosphorylatable alanines or with phospho-mimetic double glutamic acid residues (EE) ^38^ (Figure S6A). For both mutant series, we extended the number of mutant sites from either the N- or C-terminus so that different mutants covered different parts of the protein (Figure S6A). If Cdk phosphorylation stabilizes Rb, then the phospho-mutants (S/T to A) should exhibit a reduced half-life in S/G2, and the phospho-mimetic mutants (S/TP to EE) should exhibit an increased half-life in early G1. Our results are consistent with this hypothesis (Figure 3C, D; Figure S6B, C). It is worth noting that the C-terminal alanine-mutants also had a more severe cell cycle arrest phenotype (Figure S7A). This is likely because the alanine mutants do not allow the phosphorylation of C-terminal residues to disrupt Rb’s interaction with E2F-DP ^11,12,29,39^. On the other hand, the phospho- mimetic mutants did not demonstrate significant cell cycle defects (Figure S7B), likely because these Rb mutants are partially or entirely unable to bind and inhibit E2F. In addition, the introduction of phospho-mimetic mutations resulted in a smaller Rb concentration decrease in cells arrested in G1 using Palbociclib (Figure S7C, D).

Interestingly, our mutational analysis did not reveal any particular phosphorylation sites that predominantly regulated Rb’s half-life (Figure 3E, F; Figure S6C; Figure S7C, D). Instead, the degree of Rb stabilization, the ratio between early G1 and S/G2 half-lives, correlated with the total number of phospho-mimetic sites. This shows that many different phosphorylation sites contribute to Rb stability. We note that Rb14EE exhibited reduced half-lives for both early G1 and S/G2 phases, which is likely due to the additional SP230EE mutation destabilizing the protein via another mechanism (Figure 3E; Figure S7D). However, the difference between early G1 and S/G2 half-lives in Rb14EE is the smallest (Figure S6C). Collectively, these results support the hypothesis that Rb is stabilized by its hyper-phosphorylation in late G1 by Cdk (Figure 3G).

### The degradation of un- or hypo-phosphorylated Rb is mediated by the E3 ubiquitin ligase UBR5

After establishing that Rb is stabilized by phosphorylation at the G1/S transition, we next sought to identify the underlying molecular mechanism. To do this, we first tested whether Rb is degraded through the ubiquitin-proteasome system by treating cells with three commonly used inhibitors of different components of this degradation system: Bortezomib inhibits the proteasome; TAK243 inhibits the ubiquitin activating enzyme (E1); and MLN4924 inhibits the NEDD8-activating enzyme that activates the Cullin (CUL)-RING E3 ubiquitin ligases ^40,41^. We treated asynchronously growing HMEC cells with these inhibitors for 5 hours, and then immunostained the cells using antibodies for pRb (S807/811) and total Rb. TAK243 and Bortezomib treatments increased the Rb concentration in the low pRb G1 populations to a level similar to that in the high pRb G1 population, but MLN4924 did not (Figure S8A, B). RPE-1 cells (telomerase immortalized retinal pigment epithelium cells) behaved similarly to HMEC cells in that only TAK243 and Bortezomib treatments increased the Rb concentration in the low pRb G1 population (Figure S8C). To further confirm that the phosphorylation status determines Rb degradation through the ubiquitin-proteasome system, we treated cells that were induced to express un-phosphorylated Rb that lacks all the Cdk phosphorylation sites (Clover-3xFlag- RbΔCDK) or phospho-mimetic Rb (Clover-3xFlag-Rb14EE) with the three degradation inhibitors discussed above. The concentration of RbΔCDK is elevated by TAK243 and Bortezomib, but not MLN4924, and the concentration of Rb14EE does not increase following treatment by any of the three inhibitors (Figure 4A). We also confirmed the enhanced ubiquitination of RbΔCDK by pulling down Clover-3xFlag-RbΔCDK and blotting for ubiquitin. RbΔCDK was more ubiquitinated than WT Rb, which is mostly in the hyper-phosphorylated form (Figure 4B). Altogether, these data suggest that un-phosphorylated Rb is degraded in G1 through the ubiquitin-proteasome system, but not by the Cullin (CUL)-RING E3 ligases.

**Figure 4.**
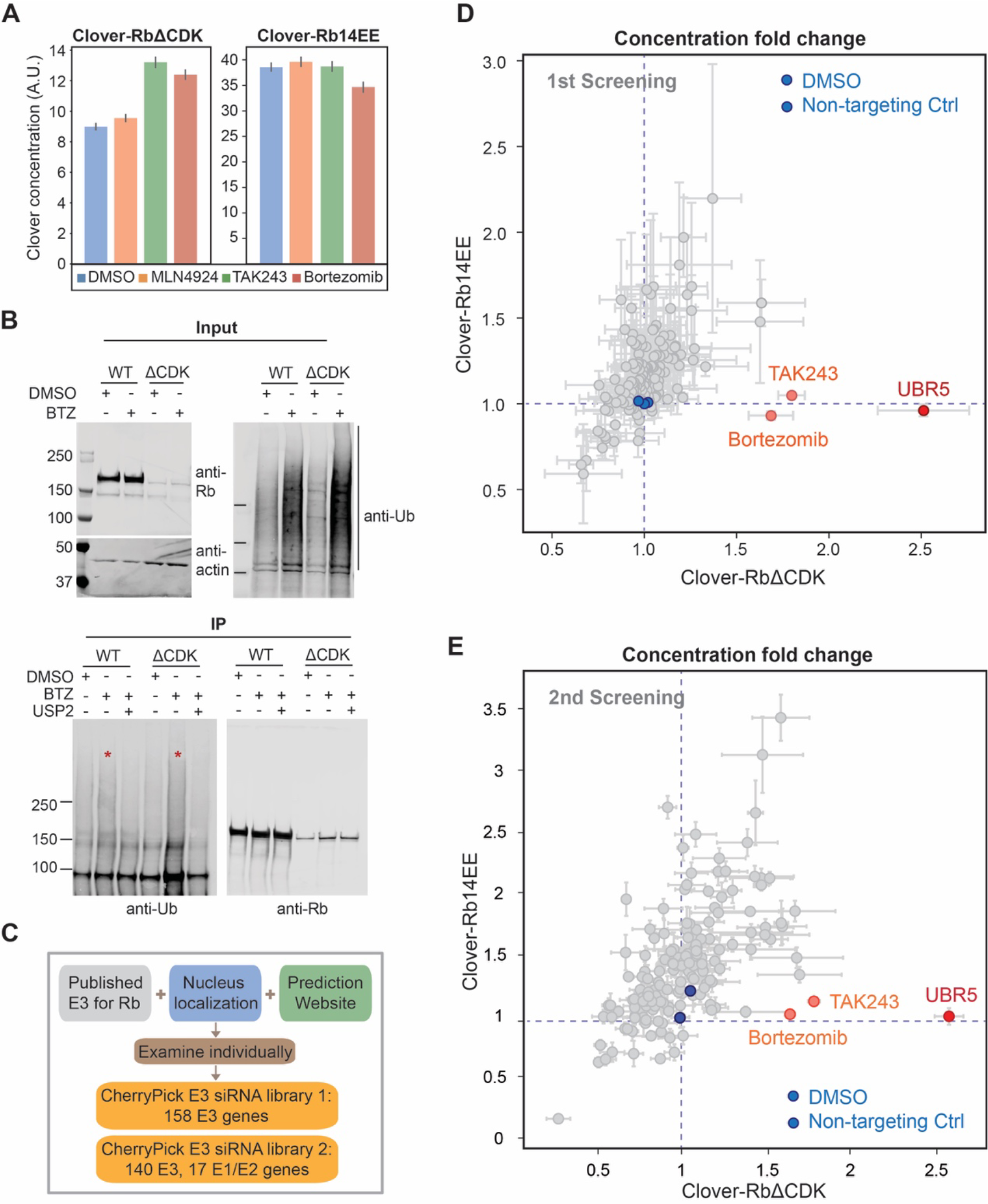
siRNA screens identified UBR5 as the E3 ligase targeting un-phosphorylated Rb for degradation A. Concentrations of Clover-3xFlag-RbΔCDK or Clover-3xFlag-Rb14EE after drug treatments. Cells expressing Clover-3xFlag-RbΔCDK and Clover-3xFlag-Rb14EE were induced with Dox (1µg/mL) for 48 hours. Then, cells were treated with the indicated drugs for 5 hours, fixed, and imaged. The concentrations of Clover-3xFlag-Rb variants were calculated by dividing total Clover intensity by nuclear area^3/2^. **B**. Un-phosphorylated Rb is degraded via ubiquitination. HMEC cells expressing Clover-3xFlag-RbΔCDK or RbWT (induced by 1µg/ml Dox for 48 hours) were treated with Bortezomib (1µM) or DMSO for 5 hours before collection. Clover-3xFlag- RbΔCDK or RbWT proteins were immunoprecipitated using anti-Flag magnetic beads. After purification, the Bortezomib treated samples were split in half and one half underwent a deubiquitination assay using USP2. The samples were then detected for Rb and Ubiquitin using immunoblot. **C.** Schematics showing siRNA library components. **D-E.** Results of the 1st and 2^nd^ siRNA screens. The concentration fold changes of Clover-3xFlag-RbΔCDK and Clover-3xFlag- Rb14EE are plotted. The fold change is calculated by dividing the Clover concentration of the treatment well by the concentration of the non-treated well. n = 4 biological replicates for the first screen and n = 3 biological replicates for the second screen.

There have been several previous studies of Rb degradation mechanisms that identified some E3 ligases ^37,42–49^. For example, MDM2 may mediate Rb degradation via its central acidic domain ^42,43,47^. The human papilloma virus (HPV) E7 protein can bind Rb and induce its degradation ^46^, which is mediated by protease cleavage at Lys810 ^44^. More recently, Cdk4/6 inhibition was found to promote Rb degradation through βTrCP1-mediated ubiquitination ^37^. To test if these E3 ligases were responsible for the observed cell cycle dynamics of Rb, we examined the effect of knocking them down on the concentration of un-phosphorylated Rb (Clover-3xFlag-RbΔCDK) and phospho-mimetic Rb (Clover-3xFlag-Rb14EE). If an E3 were responsible for Rb’s cell cycle dynamics, we would expect to see an increase in the concentration of Clover-3xFlag-RbΔCDK but not of Clover-3xFlag-Rb14EE. None of the knockdowns exhibited this predicted phenotype (Figure S9A). Even through some of the knockdowns affected the overall Rb concentration, this effect was not specific for un- phosphorylated Rb and therefore could not explain Rb’s cell cycle dynamics. Similarly, we performed the same set of knockdowns in cells arrested in G1 using Palbociclib and did not find any specific increase in the concentrations of un- or hypo-phosphorylated Rb (Figure S9B-G). This implies that there must be some additional E3 ligase responsible for the phosphorylation-dependent degradation of Rb.

To identify the E3 ligases mediating the degradation of un-phosphorylated Rb, we set up an siRNA screen. We used a customized siRNA library that included the previously published E3 ligases for Rb, some nuclear localized E3s (according to UniProt), and some additional genes predicted to be E3 ligases for Rb (http://ubibrowser.bio-it.cn/ubibrowser_v3/) (Figure 4C). HMEC cells inducibly expressing Clover-3xFlag-RbΔCDK or Clover-3xFlag-Rb14EE were transfected with the siRNA library. 48 hours later, cells were fixed and imaged. The concentration of Clover- 3xFlag-Rb variants was measured in each treatment and its fold change over non-transfected cells was calculated. As positive controls, we included the ubiquitin-proteasome system inhibitors TAK243 and Bortezomib. As expected, TAK243 and Bortezomib only increased the concentration of RbΔCDK but not Rb14EE (Figure 4D; Figure S10A, B). From this screen, we only identified UBR5 as specifically targeting un-phosphorylated Rb for degradation (Figure 4D; Figure S10A, B). UBR5 is a verified E3 ubiquitin ligase belonging to the HECT family known to play roles in transcription and the DNA damage response ^50–53^. However, Rb has never been reported to be a substrate of UBR5. To confirm that UBR5 is the main E3 ligase targeting un- phosphorylated Rb, we first performed another siRNA screen with a different siRNA library containing UBR5 and 17 other E3 genes from the first library as well as the rest of the nuclear localized E3 genes not included in the first screen. We also included several E1 and E2 genes (Figure 4C). This second siRNA screen also only identified UBR5 (Figure 4E; Figure S10B-D).

We then validated UBR5 as a hit using another two independent siRNAs against UBR5. Knockdown of UBR5 in HMEC cells led to the accumulation of un/hypo-phosphorylated Rb after Palbociclib treatment, as measured by both immunoblot and immunostaining (Figure 5A, Figure S11A). We also measured the half-life of Rb following UBR5 knockdown using live-cell imaging and found that Rb was degraded about twice as slowly in early G1, but there was no change in its stability in S/G2, as compared to the control siRNA (Figure 5B). Moreover, we examined the effect of knocking down UBR5 on HMEC cells expressing endogenously tagged Rb (*RB1- 3xFLAG-Clover-sfGFP*)^26^. Following *UBR5* knockdown, the concentration of Rb does not decrease in early G1, but is instead kept relatively constant (Figure S11B). To further confirm that the degradation of un-phosphorylated Rb by UBR5 is not cell line or cell type specific, we also examined epithelial RPE-1 cells as well as HLF (primary human lung fibroblast) and T98G (glioblastoma-derived fibroblast-like) cells. All of them showed that UBR5 knockdown increased concentrations of un- and hypo-phosphorylated Rb in Palbociclib-treated cells (Figure S11C-E).

**Figure 5.**
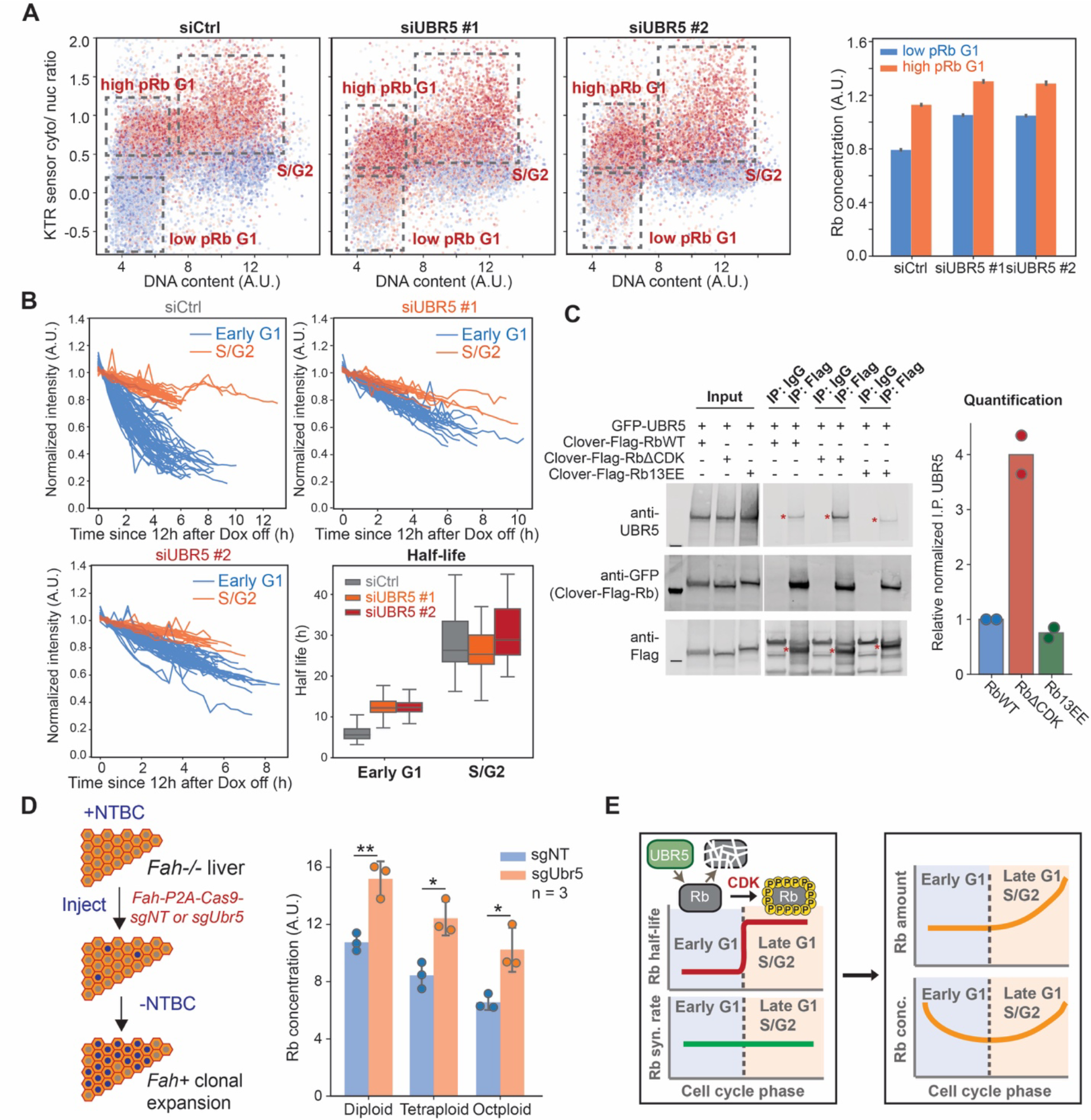
Validation of UBR5 as the E3 ligase mediating the degradation of unphosphorylated Rb A. Microscopy analysis of HMEC cells expressing the Rb (886-928) KTR sensor ^36^, which reflects Cdk activity. Cells were treated with Ctrl siRNA or *UBR5* siRNAs for 48 hours. The cytoplasmic-to-nucleus intensity ratio of the KTR sensor is plotted against DNA content (DAPI staining). The dot color indicates the Rb concentration. Quantification of the Rb concentration for different phospho-Rb levels is shown on the right. Bars indicate mean and the 95% confidence interval. **B.** Degradation traces and the calculated half-lives of Clover-3xFlag-Rb after *UBR5* knock down. Box plot indicates 5th, 25th, median, 75th and 95th percentiles. **C.** Un- phosphorylated Rb interacts with the E3 ligase UBR5. HEK293 cells were transfected with a Clover-3xFlag-RbWT, RbΔCDK, or Rb13EE construct as well as a GFP-UBR5 construct (addgene #52050). 24 hours after transfection, cells were induced by Dox. After 24 hours of induction, cells were lysed, and Flag antibodies were used to pull down Clover-3xFlag-Rb mutants. The samples were then detected for UBR5 and Rb using immunoblot. For quantification, the co-immunoprecipitated UBR5 was normalized to the immunoprecipitated Rb mutant protein amount and the input UBR5 protein amount. **D.** (Left panel) Schematic of the *Fah-/-* mouse liver model. (Right panel) Rb concentration in the low-phospho-Rb population of the primary hepatocytes isolated from mice receiving *Fah-P2A-Cas9-sgNT* or *Fah-P2A-Cas9- sgUbr5* transposons. The error bars indicate the standard deviation of the mean. **E.** Model schematic: Un-phosphorylated Rb is targeted for degradation by the E3 ligase UBR5 in early G1.

After establishing that Rb degradation depends on UBR5, we sought to test for a direct interaction between these two proteins. To do this, we performed immunoprecipitation assays with UBR5 and un-phosphorylated Rb, wild-type Rb, and phospho-mimetic Rb. We found that the un-phosphorylated Rb, which lacks all Cdk sites, can pull down significantly more UBR5 than wild-type Rb. Moreover, the phospho-mimetic Rb pulled down the smallest amount of UBR5 (Figure 5C). This result showing a direct phosphorylation-dependent interaction, is consistent with our hypothesis that UBR5 targets un-phosphorylated Rb for degradation, and Rb phosphorylation protects Rb from UBR5-dependent degradation.

To test if UBR5 mediated Rb degradation *in vivo*, we examined its effect in the mouse liver using the *Fah-/-* system^54,55^. In the *Fah-/-* system, deletion of the *Fah* gene causes toxin accumulation in hepatocytes that will lead to hepatocyte death. Toxin accumulation can be prevented by treating mice with NTBC (2-(2-nitro-4-trifluoromethylbenzoyl)-1,3- cyclohexanedione)^55^. When NTBC is withdrawn, cells expressing exogenous *Fah*, introduced by injecting *Fah+* transposons, will clonally expand to repopulate the injured liver ^55^ (Figure 5D).

Importantly, other genetic elements, such as Cas9 and gRNA, can be added to the *Fah* transposon so that they are co-integrated into some hepatocyte genomes. To knock out *Ubr5* in some hepatocytes, we modified an *Fah* transposon plasmid ^56^ and delivered *Fah-P2A-Cas9- sgUbr5* or *Fah-P2A-Cas9-sgNT* (non-targeting) transposons and the SB100 transposase into *Fah-/-* mice via hydrodynamic transfection. 8 weeks after injection, when the liver was almost fully repopulated with *Fah+* cells, we isolated the hepatocytes, plated them, and performed immunostaining or immunoblots (Figure S11F). Consistent with the results from human cell lines, knocking out *Ubr5* increased Rb concentrations in mouse hepatocytes where Rb was not hyper-phosphorylated (low pRb) (Figure 5D, Figure S11G, H). The results from this *Fah-*/- *in vivo* model further support our conclusion that the E3 ligase UBR5 targets un-phosphorylated Rb for degradation in G1 (Figure 5E).

Our results here give insight into why previous studies reported other E3s targeting Rb. First, any E3 whose knockdown results in a cell cycle phenotype would be predicted to have an effect on Rb concentration. Second, after finding Rb dynamics were driven by the degradation of un- or hypo-phosphorylated Rb in G1, we sought to find E3s that specifically targeted the un- phosphorylated RbΔCDK protein, but not the phospho-mimetic Rb14EE protein. We did find a significant increase in both RbΔCDK and Rb14EE when the E3 MDM2 was knocked down, and possibly very modest effects when other reported E3s were knocked down (Figure S9A; Figure S10A). This suggests that these other E3s might operate in different cell types or contexts, but are not generally responsible for Rb’s cell cycle dynamics.

### UBR5 and Cdk4/6 drive parallel pathways promoting the G1/S transition

It is becoming increasingly clear that there are two distinct pathways driving the G1/S transition that both operate through Rb. First, the canonical Cdk-phosphorylation pathway through which Cyclin D-Cdk4/6 complexes phosphorylate and inhibit Rb, and second, the Rb-degradation pathway that drives down the concentration of Rb in G1 phase. One prediction from this parallel pathway model is that cells lacking the Rb-degradation pathway should be more sensitive to inhibition of the remaining Rb-phosphorylation pathway. We can now test this prediction because we identified UBR5 as the E3 ligase targeting un-phosphorylated Rb in G1. To do this, we first generated clonal cell lines lacking *UBR5* from HMEC cells using CRISPR/Cas9. We randomly picked three *UBR5 WT* clones and three *UBR5 KO* clones for analysis (Figure 6A).

**Figure 6.**
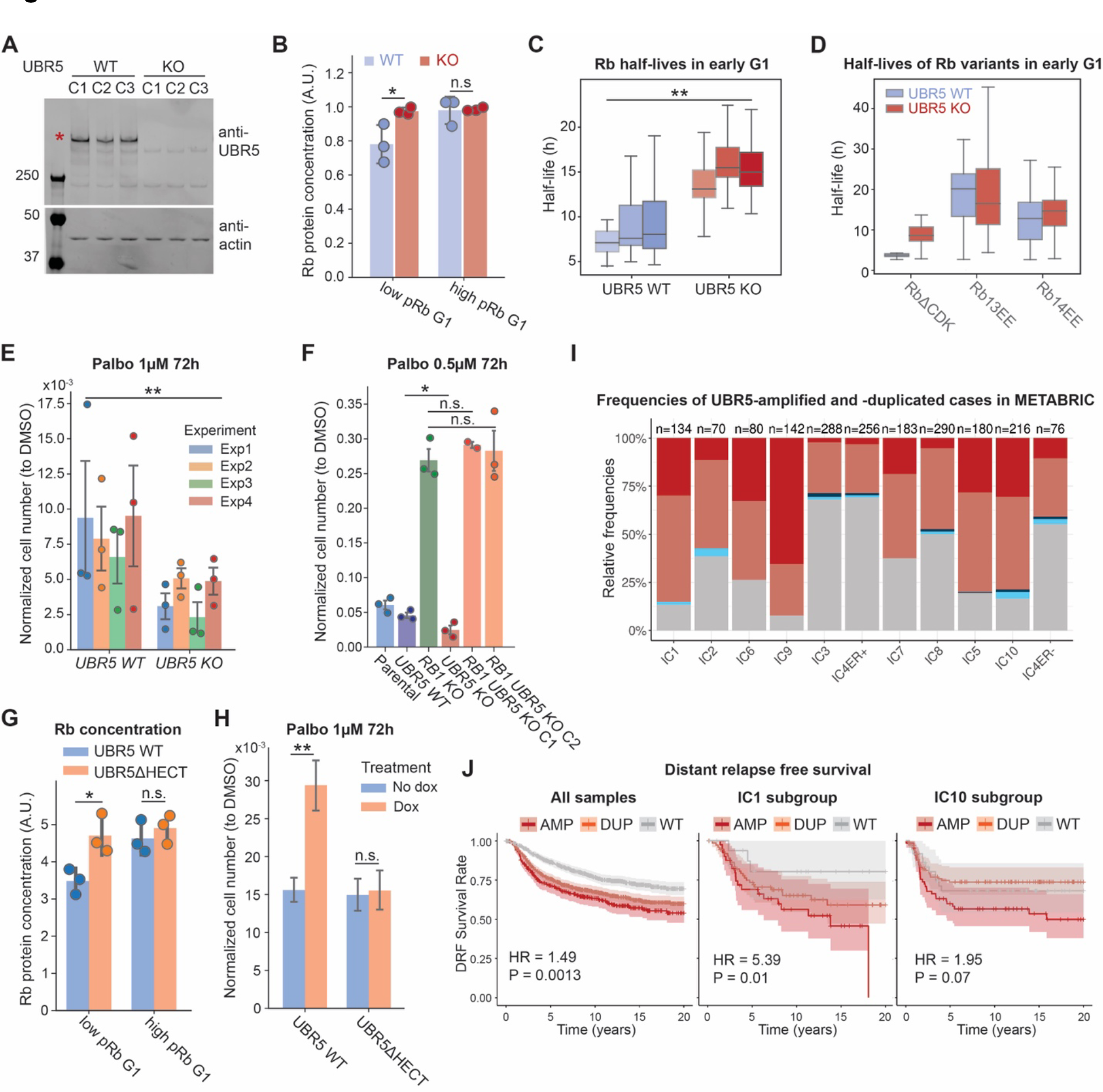
UBR5 deletion stabilizes Rb and sensitizes cells to CDK4/6 inhibition A. Immunoblot verification of *UBR5 KO* clonal cell lines. **B.** Quantification of Rb concentration in different phospho-Rb populations of *UBR5 WT* or *KO* clonal cell lines expressing the Rb (886- 928) KTR sensor ^36^. Circles denote results from individual clones and the bars denote the standard deviation. **C.** The half-life distributions of Clover-3xFlag-RbWT protein in early G1 phase from 6 clonal cell lines (3 *WT*, 3 *UBR5 KO*). **D.** The half-life distributions of Clover- 3xFlag-RbΔCDK, Rb13EE, and Rb14EE in early G1 phase in *UBR5 WT* and *UBR5 KO* cells. **E.** Normalized cell number of *UBR5 WT* and *KO* clonal cell lines after Palbociclib (1µM) treatment for 72 hours. Drug treatment started the day after plating. After 72 hours of drug treatment, cells were fixed and the cell number in each well was measured. Normalized cell number is the cell number following a Palbociclib treatment divided by the cell number following a DMSO control treatment. n = 4 biological replicates and error bars indicate the standard deviation. **F.** Normalized cell number for *UBR5 WT*, *RB1 KO*, and *RB1 UBR5* double *KO* clonal cell lines following Palbociclib (0.5µM) treatment for 72 hours. n = 3 biological replicates. **G.** Quantification of Rb concentration in different phospho-Rb populations of *UBR5 KO* cells with either *UBR5 WT* or *UBR5ΔHECT* added back by Dox-induction. Circles denote results from individual clones and the bars denote the standard deviation. **H.** Normalized cell number of *UBR5 KO* cells with *UBR5 WT* or *UBR5ΔHECT* added back following Palbociclib (1µM) treatment for 72 hours. GFP-UBR5 WT or GFP-UBR5ΔHECT were induced by Dox (100ng/ml). The results show the average from 3 *UBR5 KO* clonal cell lines with different UBR5 variants added back. We performed 3 biological replicates. The error bars indicate the standard deviation. **I.** Frequencies of *UBR5*-amplification and *UBR5*-duplication in the METABRIC dataset across the different breast cancer subtypes and IC subgroups. The number of tumors within each group is indicated on the top of each bar. AMP = amplification corresponding to a total copy number ≥ 6; DUP = duplication corresponding to a total copy number between 3 and 5; Hetero_DEL = heterozygous deletion; nLOH = neutral loss-of-heterozygosity. **J.** Kaplan-Meier curves of distant relapse-free (DRF) survival for the *UBR5*-amplified, *UBR5*-duplicated, and WT cases across all samples (left panel), in the IC1 subgroup (middle panel), and in the IC10 subgroup (right panel). The squares correspond to estimated hazard ratios and the segments correspond to their 95% confidence intervals. HR = hazard ratio. P-value tests the difference between the WT and the AMP curves corrected for the other clinical covariates. Survival analysis was performed using Cox’s proportional hazard models, comparing *UBR5*-amplified to WT cases and corrected for the clinical covariates (age, grade, tumor size, lymph node, and ER status) ^58^.

*UBR5 KO* cells exhibited higher Rb concentrations in low pRb G1 cells (Figure 6B; Figure S12A). Moreover, *UBR5 KO* cells also exhibited both higher endogenous Rb concentrations and higher exogenous Clover-3xFlag-Rb concentrations following Palbociclib treatment to arrest cells in G1 (Figure S12B, C; Figure S14A, B). *UBR5 KO* cells also exhibited increases in Rb half-life in early G1 (Figure 6C; Figure S13A, B), and exhibited increases in the half-life of un- phosphorylated RbΔCDK, but not phospho-mimetic Rb14EE, as measured by our live-cell imaging assay (Figure 6D; Figure S13C, D). These knockout lines therefore exhibited all the same effects we observed in our earlier knockdown experiments shown in Figure 5.

Having generated *UBR5 KO* cells, we are can now test the parallel pathway model prediction that cells lacking the Rb-degradation pathway are more sensitive to inhibition of the Rb-phosphorylation pathway. To do this, we treated *UBR5 WT* and *UBR5 KO* cells with the Cdk4/6 inhibitor Palbociclib for 72 hours and then measured cell proliferation by counting cell numbers. Since different clonal cell lines had different proliferation rates to begin with (Figure S15A), we normalized the cell numbers of Palbociclib treated cells to the cell numbers in the DMSO control treatment for each clonal cell line. Moreover, this normalization also accounts for the slower growth rates of *UBR5 KO* cells (Figure S15A) that are likely due to the dysregulation of other UBR5 substrates. As predicted by the parallel pathway model, *UBR5 KO* cells are more sensitive to Palbociclib treatment than *UBR5 WT* cells (Figure 6E, Figure S15A). To further determine the proliferation status of *UBR5 WT* and *KO* cells, we also stained the cells with phospho-Rb antibodies following prolonged DMSO or Palbociclib treatments up to 6 days. As expected, a significantly higher proportion of *UBR5 WT* cells were progressing through the cell cycle (as indicated by cells having hyper-phosphorylated Rb) compared to *UBR5 KO* cells, again supporting the parallel pathway model (Figure S15B-E).

Since UBR5 has other substrates that may also affect cell cycle progression, we wanted to examine if UBR5 KO cells’ increased sensitivity to Palbociclib treatment was due to the stabilization of Rb. To test this, we knocked out *RB1* in *UBR5 KO* cells using CRISPR/Cas9 (Figure S16A) and tested their sensitivity to Palbociclib treatment. Knocking out *RB1* in *UBR5 KO* cells completely rescued the increased Palbociclib sensitivity exhibited by *UBR5 KO* cells (Figure 6F; Figure S16B), suggesting that the effect of UBR5 on the G1/S transition is primarily through the stabilization of Rb. Lastly, to make sure that the effect of UBR5 on cell cycle progression was due to its E3 ligase activity, we added back either wild-type UBR5 or an inactive mutant UBR5 to the *UBR5 KO* cells using the Dox-inducible system (Figure S17A). The mutant UBR5 has a C2768A mutation in the HECT domain (abbreviated as UBR5ΔHECT), which kills its catalytic activity ^51^. Consistent with the role of UBR5 degrading un-phosphorylated Rb, the expression of UBR5 WT was able to reduce Rb concentration in the low pRb G1 population in *UBR5 KO* cells, but expressing UBR5ΔHECT did not (Figure 6G; Figure S17B, C). We treated cells expressing UBR5 WT or UBR5ΔHECT (induced by Dox) with DMSO or Palbociclib for 72 hours, and found that adding back UBR5 WT decreased the cells’ Palbociclib sensitivity compared to the no Dox control, whereas adding back the UBR5ΔHECT did not (Figure 6H; Figure S18A, B). This indicates that the E3 ligase activity of UBR5 is essential for its function as a key component of the Rb-degradation pathway.

Since deleting UBR5 sensitizes cells to treatment by Cdk4/6 inhibitors that are currently used to target breast cancers, we next explored if UBR5 itself could be a potential therapeutics target. To do this, we examined the *UBR5* copy number level in breast cancer patients and how it relates to the patient’s survival. Since there is a lot of variability between patients depending on the molecular basis of their disease, we categorized patients for analysis using the Integrative Cluster subtype (IC) framework that partitions groups of patients based on multiple types of genomic data ^57,58^. IC subtypes predict patient outcomes beyond the historical clinical stratification based on hormone receptors status: ER+, HER2+, and triple-negative subgroups ^57,58^. IC classification stratifies ER+ tumors into ER+ Typical-risk (IC3,4ER+,7,8) and ER+ High-risk of relapse (IC1,2,6,9) categories, and stratifies triple-negative tumors (TNBC) into the genomically stable and unstable subtypes IC4ER- and IC10, respectively ^57,57^. Among the 1894 patient samples from the Molecular Taxonomy of Breast Cancer International Consortium (METABRIC), the *UBR5* gene was amplified in the majority of the patients. When stratified by the prognostic subtypes,

*UBR5* gene copy amplification was observed in all subgroups but to various degrees (Figure 6I; Figure S18C). Importantly, *UBR5* amplification was particularly dominant in subgroups associated with a worse prognosis (ER+ High-risk, HER2+, and TNBC tumors). We then assessed how *UBR5* gene copy alteration affected patient survival. By analyzing the distant relapse-free (DRF) survival of these patients, we found that *UBR5* amplification is associated with worse DRF survival (Figure 6J), especially in IC1 (ER+ High-risk) and IC10 (TNBC, basal-like tumors) subtypes (Figure 6J). These results suggest that breast cancer cells might increase *UBR5* expression through copy number amplification to facilitate cell cycle progression.

## Discussion

Two distinct pathways drive the G1/S transition by reducing the activity and amount of the key cell cycle inhibitor Rb: the canonical Cdk-phosphorylation pathway and the Rb-degradation pathway (Figure 7). Here, we report that the Rb-degradation pathway is driven by the E3 ubiquitin ligase UBR5 and is shut off by Rb hyper-phosphorylation in late G1. This mechanism could explain why some cells can still progress into S phase and activate Cdk2 even when Cdk4/6 is being inhibited ^7^. Namely, when Cdk4/6 activity is inhibited in cells, protein degradation continuously decreases the Rb concentration. This eventually leads to partial activation of E2F-dependent transcription, which activates Cyclin E/A-Cdk2 to hyper- phosphorylate Rb and drive S phase entry. Therefore, the Rb degradation pathway provides cells with an alternative path to enter the division cycle, emphasizing the plasticity of the cell cycle program that is likely important in different cell types and contexts.

**Figure 7.**
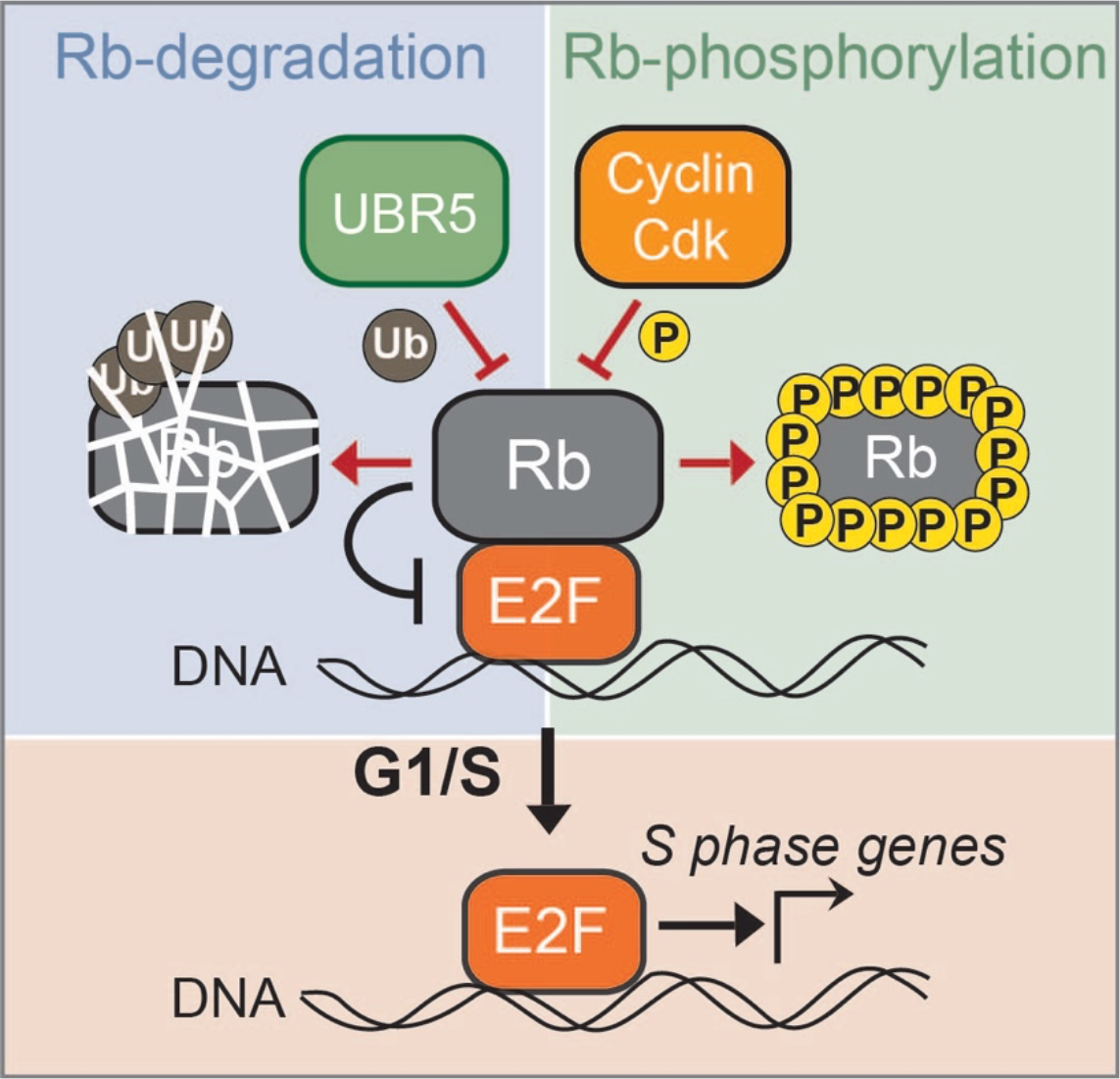
Model schematic of parallel pathways regulating Rb activity and thereby the G1/S transition. Rb is targeted for degradation by UBR5 in parallel to the canonical Rb phosphorylation pathway.

While the two pathways impinging on Rb activity employ distinct mechanisms, degradation and phosphorylation, their activities become interconnected once cells begin the G1/S transition. Namely, the degradation of Rb eventually drives an increase in E2F activity and thereby transcription of Cyclin A and Cyclin E. These downstream cyclins can then form a complex with Cdk2 to drive Rb hyper-phosphorylation and stabilization. Consequently, Rb degradation drives Cdk activity, which in turn inhibits Rb degradation as cells progress into S phase. Nevertheless, while these activities become interconnected at the restriction point and the G1/S transition, prior to this point it is likely that they operate independently so that UBR5 and Cyclin D-Cdk4/6 activities can be separately modulated to control the timing of the G1/S transition in a variety of cellular contexts.

The parallel pathway model predicts that cells with low Cdk activity will more heavily rely on the Rb-degradation pathway to facilitate their G1/S transition. This prediction is supported by our live-cell imaging results (Figure 1). As reported previously, we see many cultured cells immediately activating Cdk activity (Cdk-high), while some other cells have a longer G1 period with lower Cdk activity (Cdk-low) ^1,25^. In Cdk-low cells, the Rb concentration continually decreases in G1 until a couple of hours before S phase. Moreover, the Rb concentration in G1 negatively correlates with the G1/S transition rate. While manipulating Rb concentration has minor effects on cell proliferation in many cancer cell lines and immortalized cell lines ^16,32^, this is likely due to the high Cdk activity in these cells and the rich *in vitro* culture environment.

Consistent with this view, Rb concentration is not important in Cdk-high activity cells, which are the majority in most *in vitro* cultured cell lines. The cell cycle dynamics of cultured cells stands in sharp contrast to the dynamics of cells proliferating *in vivo*. For example, the epidermal stem cells in mouse skin have a G1 length around 48-72 hours^28,33^, far exceeding the G1 length in *in vitro* cultured cell lines ^1,2,25^. Thus, cells proliferating *in vivo* are likely more similar to the Cdk-low cells and therefore are expected to rely on the Rb degradation pathway. Indeed, manipulation of the Rb-pathway in mouse epidermal stem cells regulates their G1 dynamics^28^.

While we identified UBR5 as the E3 ligase that targets Rb for degradation, we want to emphasize that Rb is not the sole target of UBR5. Multiple papers have shown that UBR5 has other targets including MYC, Nuclear hormone receptors, and SPT4/5 ^51–53^. Here, we identify Rb as an additional target of UBR5. Yet, the mechanism through which UBR5 recognizes Rb is still unclear. UBR5 likely engages its substrates as a dimer or tetramer, which can target distinct degron linear motifs as indicated by recent cryo-EM structures ^52,53,59–61^. Since UBR5 is a large multi-domain protein, it is possible that different docking mechanisms can be utilized for engaging different groups of substrates ^51,53,60–62^. Interestingly, two recent studies proposed that UBR5 targets its substrates on chromatin ^52,53^. This possible preference of UBR5 for chromatin-bound targets might explain the results of our mutational analysis of Rb. Namely, the more tightly an Rb variant is predicted to bind the E2F transcription factor, the more rapidly it is degraded. The Rb variants with increasing numbers of phospho-mimetic sites (Figure 3E; Figure S6A, C) have decreasing binding affinity to E2F (and chromatin), and exhibited increasing stability in G1. In support of such a model, the stabilization of Rb at the G1/S transition is coincident with its hyper-phosphorylation and dissociation from the chromatin-bound E2F transcription factors ^63^.

UBR5, like other components of the canonical Rb-phosphorylation pathway, appears frequently mutated in cancer and may become a target for therapies ^64^. For example, Cdk4/6 inhibitors in combination with endocrine therapy are used to treat advanced estrogen receptor positive (ER+)/human epidermal growth factor receptor-2 negative (HER2-) breast cancers ^65–67^.

However, this application is frequently limited by the intrinsic and acquired therapeutic resistance observed in patients ^68,69^. One possible way to improve upon current therapies targeting the Rb-phosphorylation pathway is to also target the Rb-degradation pathway through UBR5. Since deleting UBR5 sensitizes cells to treatment by Cdk4/6 inhibitors, it is possible that current Cdk4/6 inhibitor-based treatments for breast cancer can be improved by developing novel therapeutics targeting the Rb-degradation pathway through UBR5, which, intriguingly, is frequently amplified in such cancers (Figure 6I).

The inability of current therapies to inhibit cell division reflects our incomplete knowledge of the signaling pathways involved. During G1 phase, the cell integrates many signals to make the decision to commit to cell division including two separate signaling pathways targeting Rb. The existence of these two parallel pathways explains the ability of some cells to proliferate in the absence of Cyclin D-dependent kinase activity ^7,70,71^. Namely, these cells rely more heavily on the UBR5-Rb degradation pathway than the canonical Cdk4/6 Rb-phosphorylation pathway. In addition to providing such a robust entry to the cell division cycle in a particular cellular context, the existence of parallel pathways regulating the G1/S transition might be due to the different proliferative requirements of diverse cell types ^72^. Different pathways can be independently tuned to precisely calibrate the rate of proliferation required by the myriad cell types and niches of a multicellular organism. Discovering the mechanisms underlying these G1/S regulatory pathways, such as we have here for Rb-degradation, will therefore give insight into both development and disease.

## Acknowledgements

We thank Julien Sage, Seth Rubin, Hao Zhu, Peter Pryciak and members of the Skotheim laboratory for discussions and feedback on the manuscript. We thank Hao Zhu for sharing the Fah transposon plasmids. We thank Markus Grompe for sharing the Fah-/- mice. This work was supported by a Chan Zuckerberg Biohub Investigator Award (J.M.S.), and the NIH (P01 CA254867 and K99GM147351). The siRNA screens were done using the ImageXpress Micro CONFOCAL microscope at the High Throughput Screening Knowledge Center (HTSKC) at Stanford, which was funded by the NIH Shared Instrumentation Grant (S10OD026899).

## Author contributions

S.Z. performed the experiments, L.V. performed the mathematical modeling, E.Z., S.Z, and J.M.S. designed the experiments, L.M. and C.C performed the cancer genomics analysis, and S.Z. and J.M.S. wrote the manuscript.

## Materials and Methods

### Mice

All mice were handled in accordance with the guidelines of the Institutional Animal Care and Use Committee at Stanford University. The *Fah^-/-^* mice were kindly shared by Dr. Markus Grompe’s group at Oregon Health & Science University. At 6 weeks old, the *Fah^-/-^* mice were subjected to hydrodynamical transfection through their tail vein. We injected one plasmid containing a transposon that carries *Fah*, *Cas9*, and a guide RNA, and a second plasmid containing the transposase SB100. The plasmid backbone was kindly provided by Dr. Hao Zhu’s lab at UT Southwestern Medical Center. We modified the guide RNA sequence on the transposons to either a negative control guide RNA sequence or a guide RNA targeting the *Ubr5* gene. 8 weeks after the injection, the mice were sacrificed to isolate primary hepatocytes.

### Cell culture conditions and cell lines

All cells were cultured at 37°C with 5% CO2. Non-transformed hTERT1-immortalized human mammary epithelium cells (HMEC) were obtained from Stephen Elledge’s laboratory at Harvard Medical School^73^ and cultured in MEGM™ Mammary Epithelial Cell Growth Medium (Lonza CC- 3150). In microscopy experiments we used the same media but without phenol red to reduce background fluorescence (Lonza CC-3153 phenol-red free basal media supplemented with growth factors and other components from the Lonza CC4136 kit). T98G cells were purchased from ATCC, recently isolated primary fetal human lung fibroblasts (HLF) were purchased from Cell Applications, and hTERT1-immortalized retinal pigment epithelium (RPE-1) cells were obtained from the Cyert laboratory at Stanford. All these cell lines were grown in Dulbecco’s modification of Eagle’s medium (DMEM) with L-glutamine, 4.5 g/l glucose, and sodium pyruvate (Corning), supplemented with 10% FBS (Corning) and 1% penicillin/streptomycin.

### Fluorescent reporter cell lines

The S/G2 component of the FUCCI cell cycle reporter – mCherry-Geminin – was cloned into the CSII-EF-MCS lentiviral vector backbone under a constitutive EF1α promoter^1^. The CSII vector, the lentiviral packaging vector dr8.74, and the envelope vector VSVg were transfected into HEK 293T cells by PEI (1mg/mL, Sigma-Aldrich). 48 hours later, the lentivirus-containing medium was collected and used to infect HMEC cells. 2-3 days after infection, positive cells were sorted by FACS and expanded. The endogenously tagged *RB1-3xFLAG-Clover-sfGFP* HMEC cell line was created by Evgeny Zatulovskiy^26^.

### Inducible expression of wild-type and mutant Rb in cell lines

To inducibly express wild-type Rb and Rb mutants in cells, we used the doxycycline-inducible Rb cassette published in^29^, and performed site-directed mutagenesis (E0554S, NEB) to generate Rb mutant plasmids. All the plasmids contained the wild-type *RB1* gene or *RB1* mutants fused with fluorescent Clover and 3xFLAG affinity tag sequences, a zeocin resistance gene, and a Tet-On 3G transactivator gene driven by the EF1α promoter. The HMEC cell lines stably expressing doxycycline-inducible Rb variants were generated by transfecting cells with 1 µg of doxycycline- inducible plasmid and 1 µg of PiggyBac Transposase plasmid using the FuGene HD reagent (Promega E2311). Zeocin (300 µg/ml) selection began two days after transfection and lasted for at least 2 weeks until all the cells became resistant.

### *UBR5* knockout cell lines and *RB1* knockout cell lines

*UBR5* knockout HMEC cells and *RB1* knockout HMEC cells were generated using the CRISPR knockout kit v2 from Synthego following the manufacturer’s protocol. Briefly, the Ribonucleoprotein (RNP) complexes (9:1 sgRNA to Cas9 ratio) were assembled in Nucleofector Solution plus Supplement (Lonza Amaxa HMEC Nucleofector Kit) to a total volume of 100µL. The RNP was incubated at room temperature for 10 minutes. 0.5 million HMEC cells were resuspended in 100µL RNP solution and transferred to a cuvette. This was followed by electroporation using the Lonza nucleofector 2b device (program Y-001). 48 hours later, the knockout efficiency was examined by western blot. In both cases, we obtained about 40-50% knockout efficiency. The resulting cell populations were then single cell sorted to make clonal cell lines. After single cell expansion, the grown out clones were validated for *RB1* or *UBR5* knockout, and wild-type clones and knockout clones were kept for further analysis. The *UBR5 RB1* double knockout cell lines were generated by knocking out the *RB1* gene in the *UBR5* knockout clones.

### Primary hepatocyte isolation and 2D culture

Primary hepatocytes were isolated by two-step collagenase perfusion^74^ using liver perfusion medium (Thermo Fisher Scientific, 17701038), liver digest medium (Thermo Fisher Scientific, 17703034), and hepatocyte wash medium (Thermo Fisher Scientific, 17704024). The protocol for 2D culture of primary hepatocytes was kindly shared by Yinhua Jin from Dr. Roeland Nusse’s lab at Stanford University^75^. Briefly, primary hepatocytes from *Fah-/-* mice were isolated by two-step collagenase perfusion. After isolation, cells were washed 3 times with hepatocyte wash medium (Thermo Fisher Scientific, 17704024). Cells were then plated in a 6-well plate precoated with collagen I (50µg/mL) at a density of 200,000 cells per well. The culture medium contained 3µM CHIR99021 (Peprotech), 25ng/mL EGF (Peprotech), 50ng/mL HGF (Peprotech), and 100ng/mL TNFa (Peprotech) in Basal medium. The Basal medium contained William’s E medium (GIBCO), 1% Glutamax (GIBCO), 1% Non-Essential Amino Acids (GIBCO), 1% Penicillin/streptomycin (GIBCO), 0.2% normocin (Invitrogen), 2% B27 (GIBCO), 1% N2 supplement (GIBCO), 2% FBS (Corning), 10mM nicotinamide (Sigma), 1.25mM N-acetylcysteine (Sigma), 10µM Y27632 (Peprotech), and 1µM A83-01 (Tocris). The culture medium was refreshed every other day. Cells were passaged via trypsinization using TrypLE (Thermo-Fisher). The first passage cells were used for immunostaining or western blot analysis.

### Live cell imaging and analysis

The cells for imaging were seeded on 35-mm glass-bottom dishes (MatTek) one day before imaging. Then, the cells were transferred to a Zeiss Axio Observer Z1 microscope equipped with an incubation chamber and imaged for 48 hours at 37°C and 5% CO2. Brightfield and fluorescence images were collected at multiple positions every 20 minutes using an automated stage controlled by the Micro-Manager software. We used a Zyla 5.5 sCMOS camera and an A-plan 10x/0.25NA Ph1 objective. For half-life measurements, cells were plated in doxycycline containing medium (1µg/mL) and induced for 36 hours. After this induction, the doxycycline-containing medium was removed, the cells were washed once with fresh doxycycline-free medium, and then, once the fresh medium lacking doxycycline was added, the cells were transferred to the microscope for imaging. For most of the Rb mutants, the half-life analysis started 12 hours after doxycycline removal to eliminate the potential effect of protein synthesis from residual mRNA. For the fast degraded Rb mutants, e.g., RbΔCDK, the half-life analysis started 3 hours after doxycycline removal to ensure the signal-to-noise ratio is good enough for accurate quantification. The cell cycle stage was classified using an mCherry-Geminin FUCCI sensor. The early G1 phase traces were taken as those having no mCherry-Geminin expression that lasted longer than 7 hours. The S/G2 phase is defined by mCherry-Geminin FUCCI marker expression. For cells expressing the HDHB CDK sensor^25^, the transition from low CDK activity to high CDK activity was taken as the inflection point of the cytoplasm-to-nuclear fluorescence ratio ^25^. The volume of cell nucleus was used as a proxy of total cell volume because nuclear volume is known to scale in proportion to cell volume and the the nucleus can be segmented and measured much more accurately that the irregular-shaped cell^76^.

### siRNA transfection

For siRNA screening, the library was purchased from Horizon Discovery. We constructed two customized libraries against the target genes listed in Extended Data Table 1. We used ON- TARGETplus siRNA (Horizon Discovery), with 4 different siRNA sequences targeting the same gene pooled together. siRNA transfection was performed using Lipofectamine RNAiMAX (Invitrogen) following the manufacturer’s protocol. Briefly, for reverse transfection, 7500 HMEC cells with Clover-3xFlag-Rb variant cassettes (15000 cells if the variant is RbΔCDK because the cells will be arrested in G1) suspended in 100µL doxycycline containing medium (1µg/mL) were added per well into a 96-well plate containing a mixture of pooled siRNAs (1.5pmol per well) and Lipofectamine RNAiMAX (0.2µL per well in 20µL OptiMEM). Cells were subsequently grown for 48 hours at 37°C before fixation. For MLN4924, TAK243, and Bortezomib treatments, the drugs were added 5 hours before fixation. Cells were fixed with 4% paraformaldehyde for 20 minutes at room temperature, permeabilized with 0.5% Triton™ X-100 (Sigma-Aldrich) for 15 minutes at room temperature, and then incubated with 500 nM DAPI for 30 minutes at room temperature. After PBS wash, cells were imaged using the ImageXpress Micro Confocal at the High- Throughput Screening Knowledge Center of Stanford.

For regular siRNA knockdown, we used the Silencer-Select Pre-designed siRNA (Ambion Thermo Fisher). For most genes, we purchased 2 different siRNAs. Lipofectamine RNAiMAX (Invitrogen) was used for siRNA transfection. For 24-well plates, cells were plated 1 day before such that they were ∼40% confluent at the time of transfection. For each well, 6 pmol of siRNA in 50 µl Opti- MEM was mixed with 1 µl RNAiMAX in 50 µl Opti-MEM. After 10-20 minutes of incubation at room temperature, the mixture was added to the cells. 48 hours later, the cells were lysed for western blot or qPCR analysis.

### Palbociclib sensitivity assay

Cells were plated in 96-well plates (2000/well) 1 day before drug treatment. For cells with inducible Rb cassettes, cells were plated in different concentrations of doxycycline (0, 20ng/ml, 50ng/ml, 150ng/ml, 500ng/ml, 1000ng/ml) to induce different concentrations of Clover-3xFlag-Rb. Palbociclib or DMSO were added the next day, and the medium was refreshed every day. After 3 days of drug treatment, cells were fixed with 4% paraformaldehyde for 20 minutes at room temperature, permeabilized with 0.5% Triton™ X-100 (Sigma-Aldrich) for 15 minutes at room temperature, and then incubated with 500 nM DAPI for 30 minutes at room temperature before imaging. After PBS wash, cells were imaged using the ImageXpress Micro Confocal at the High- Throughput Screening Knowledge Center of Stanford. Cell nuclei were segmented and counted to indicate cell number. 4 technical replicates were prepared for each condition (4 wells for the same genotype and same treatment), and the average was used for the final cell count.

### Immunofluorescence staining

Cells were seeded on a 35-mm glass-bottom dish (MatTek) or a 6/24-well glass-bottom plate (Cellvis) one day before immunofluorescence staining. For this staining, cells were fixed with 4% paraformaldehyde for 20 minutes at room temperature, permeabilized with 0.5% Triton X-100 (Sigma-Aldrich) for 15 minutes at room temperature, and then blocked with 3% BSA in PBS. Then, the cells were incubated with primary antibodies overnight at 4°C. After 3 washes with PBS, the cells were incubated with Alexa Fluor secondary antibodies (Invitrogen, A32728) at 1:1000 for 1 hour at room temperature. After 3 washes with PBS, cells were incubated with 500 nM DAPI for

30 minutes at room temperature before imaging. The primary antibodies used for immunofluorescence were Anti-Rb (Santa Cruz, sc-74570, 1:100), Anti-phospho-Rb (S807/811) (CST, 8516, 1:400). The cells were imaged using a Zeiss Axio Observer Z1 microscope with an A-plan 10x/0.25NA objective. For the 24-well plates, cells were imaged using the ImageXpress Micro Confocal from the High-Throughput Screening Knowledge Center at Stanford.

### Flow cytometry and cell sorting

For flow cytometry analysis, cells were grown on 6-well plates to ∼70% confluence and harvested following trypsinization. The cells were then fixed with 4% formaldehyde for 10 minutes at 37°C and permeabilized with 90% ice-cold methanol for 30 minutes on ice. Fixed and permeabilized cells were washed once with PBS, blocked with 3% BSA in PBS for 30 minutes at 37°C, and then stained with primary antibodies for 2 hours at 37°C. The cells were then washed twice with a wash buffer (1% BSA in PBS + 0.05% Tween® 20), stained with the fluorophore-conjugated secondary antibodies (Life Technologies) at 1:1000 dilution for 1 hour at 37°C, and then washed twice again. After this treatment, the cells were resuspended in PBS containing 3 µM DAPI for DNA staining, incubated for 30 minutes at room temperature, and then analyzed on an Attune NxT Flow Cytometer (Thermo Fisher Scientific). For live cell flow cytometry analysis, cells were harvested from dishes by trypsinization, stained with 20 µM Hoechst 33342 DNA dye in PBS for 30 minutes at 37°C and then analyzed on an Attune NxT Flow Cytometer (Thermo Fisher Scientific). For live cell sorting, the cells were harvested from dishes following trypsinization, resuspended in fresh medium, stained with Hoechst (if sorting by cell cycle phase), and then sorted on a BD FACSAria flow cytometer. DNA content and the mCherry-Geminin fluorescent reporter were used to determine cell cycle phase, and the side scatter area parameter was used as a readout for cell size. During sorting, cell samples were kept at 4°C, and the sorted cells were collected for further RNA isolation and RT-qPCR analysis or for immunoblotting. For single clone derivation, cells were sorted into 48-well plates containing growth medium using the single-cell sorting mode.

### RNA immunoprecipitation assay

We performed RNA immunoprecipitation following published protocols ^77^. Briefly, cells were lysed in an ice cold polysome lysis buffer with 3-5 million cells per 1 ml of lysis buffer. The lysate was spun down at 16000xg for 15 minutes at 4°C, and the supernatant was transferred to a new tube. 50µl of equilibrated protein A/G-agaraose beads (Pierce, Thermo Fisher) were added to the supernatant for preclearing at 4°C for 1h. 100µl of the lysate was saved as the input sample, and the rest was incubated with 5µg of eIF4E antibody (sc-271480, Santa Cruz) or Mouse IgG (sc- 2025, Santa Cruz) at 4°C overnight. The next day, 50µl of equilibrated protein A/G-agaraose beads were added to each sample and rotated at 4°C for 4 hours. Then, 100µl of the supernatant was saved for the flow-through sample, and the beads were washed with lysis buffer 2 times, and then washed with lysis buffer containing 1M urea another 2 times. Then, the beads were boiled in Tris-EDTA containing 1% SDS and 12% β-Mercaptoethanol before RNA was extracted using a Direct-zol RNA Microprep kit (Zymo Research). The extracted RNA was then prepared for RT- qPCR analysis to examine *RB1, Actin*, and *GAPDH* expression.

### Co-immunoprecipitation

Cells were lysed in lysed in 1% NP-40 lysis buffer (50 mM HEPES, 150 mM NaCl, 1% NP-40, 1 mM EDTA with protease inhibitor mixture and 1 mM PMSF). Lysates were incubated on ice for 30 minutes before clearing by centrifuging at 16,000 × g at 4°C for 20 min. Protein lysates were pre-cleared with Protein A/G plus agarose beads (SCBT, sc-2003) at 4°C for 1 hour. Then, 8% of cleared lysates were taken as input, the rest were incubated with either 5 µg FLAG M2 antibody (Millipore Sigma #F1804) or Mouse IgG control (BioLegend, 400101) on a rotor at 4°C for 2 hours. Then, Protein A/G plus agarose beads (SCBT, sc-2003) beads were added to each sample, incubated on a rotor at 4°C for 2 hours. Then, beads were washed in 1% NP-40 lysis buffer 3 times, 10 minutes each time at 4°C on a rotor. The immunoprecipitated proteins were then eluted in 1x Sample Buffer 2 (Invitrogen, 1981103) by boiling at 95°C for 10 minutes. And then, the samples were analyzed by immunoblot.

### Immunoblots

Cells were directly lysed with 1x NuPAGE LDS Sample buffer (Invitrogen), and then incubated at 95°C for 10 minutes. Lysates were separated on NuPAGE 3-8% Tris-Acetate protein gels (Thermo Fisher Scientific), and transferred to nitrocellulose membranes. Membranes were then blocked with SuperBlock™ (TBS) Blocking Buffer (Thermo Fisher Scientific) and incubated overnight at 4°C with primary antibodies in 3% BSA solution in PBS. The primary antibodies were detected using the fluorescently labeled secondary antibodies IRDye® 680LT Goat anti-Mouse IgG (LI-COR 926-68020) and IRDye® 800CW Goat anti-Rabbit IgG (LI-COR 926-32211).

Membranes were imaged on a LI-COR Odyssey CLx and analyzed with LI-COR Image Studio software. The following primary antibodies were used: anti-β-Actin (Sigma, A2103, 1:2000), anti- Rb (Santa Cruz, sc-74570, 1:500), anti-UBR5 (ABE2863, Millipore, 1:2000), anti-Tubulin (T5168, Sigma, 1:2000), and anti-GAPDH (AB2302, Sigma, 1:2000).

### Image analysis

For live-cell imaging microscopy data, cell nuclei were segmented using the GFP channel from either the endogenously expressed Rb-3xFLAG-Clover-sfGFP or the inducibly overexpressed Clover-3xFlag-Rb variants. For fixed cell imaging microscopy data, cell nuclei were segmented using the DAPI DNA staining signal. Segmentation was performed using the Fiji plugin StarDist, which is a deep-learning tool for segmenting nuclei in images that are difficult to segment using thresholding-based methods. The total pixel intensities within the segmented masks in each channel were recorded, and each object’s background was subtracted based on the median intensity of the image. Nuclear volume was used as a proxy for cell size, and calculated as the nuclear area^3/2^. The tracking of live cells was done manually using the TrackMate plugin in Fiji. To determine the protein half-life, the degradation traces were fitted with an exponential decay function N = N0exp(-kt) so that the half-life t1/2 = (1/k)(ln2).

### RNA extraction and RT-qPCR

Total RNA was isolated using a Direct-zol RNA Miniprep kit (Zymo Research). For RT-qPCR, cDNA synthesis was performed with 1μg of total RNA using an iScript Reverse Transcription Kit (Biorad). qPCR reactions were made with the 2x SYBR Green Master Mix (Biorad). Gene expression levels were measured using the ΔΔCt method.

### Mathematical Model of Rb concentration dynamics

The mathematical model of Rb dynamics consists of two main equations. The first describes the time evolution of cell mass as 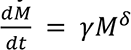, where *M* denotes the cell mass, δ<1 quantifies the departure from pure exponential growth as measured in ^78^, and γ is a constant. The second equation describes the evolution of the amount of Rb in the cell as 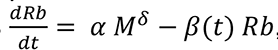, where α is an effective synthesis rate and *β*(*t*) is a time-varying degradation rate. Therefore, the amount of Rb synthesized at a given time is proportional to the cell’s growth rate. The distinctive element in this model is the time-dependence of the degradation rate *β*(*t*), which reflects the cell-cycle- dependent changes in Rb stability. It is encoded as a step change such that *β*(*t*) = *β*_’_as long as the cell is in G1 phase and that *β*(*t*) = ε*β*_0_, ε < 1, when the cell is in S/G2/M phase. To compare the relative contributions of cell-cycle-dependent synthesis and degradation changes, we also consider the case where *β* is a constant and α increases slightly in a step change at the G1/S transition. The cell goes through the G1/S transition when the concentration of Rb in the cell crosses a threshold value set *a priori*. The S/G2/M phase is represented as a timer in the model. The model is integrated in time with an Euler forward method. From arbitrary initial conditions, the model is run through a “burn-in” period until it reaches a limit cycle described by the input parameters. The model is available on GitHub (https://github.com/LucasFuentesValenzuela/RB_model).

### Genomic analysis of breast cancer patients

We extracted the copy number data from Pereira et al., Nat. Commun 2016 ^79^ (ASCAT) and we used the IC subtypes reported in the Rueda et al. Nature 2019 study ^58^. We performed survival analysis using Cox’s proportional hazard models using R package survival (version 3.5-7) and corrected for key clinical covariates as described in ^58^ (*e.g.*, age, grade, tumor size, lymph node, and ER status). We generated Kaplan–Meier and forest plots using the R package survminer (version 0.4.9).

### Statistical analysis

The data in most Fig. panels reflect multiple biological replicate experiments performed on different days. The mouse experiments used mice derived from different litters. For comparison between groups, we generally conducted unpaired two-tailed Student’s t-tests. Statistical significance is displayed as p < 0.05 (∗) or p < 0.01 (∗∗) unless specified otherwise.

---

**Figure S1.**
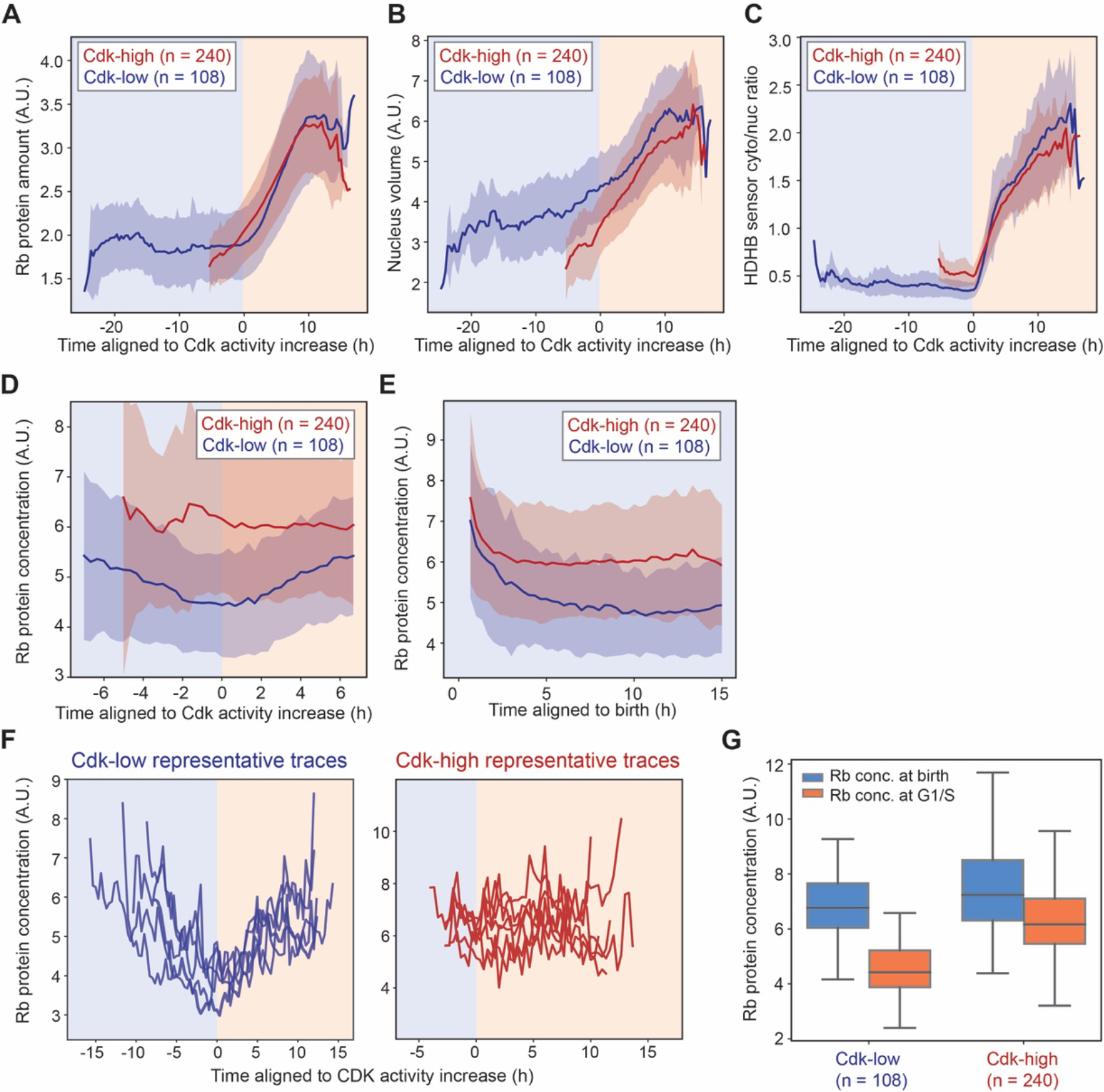
Rb concentration continuously decreases in early G1 phase in Cdk-low cells. A-E. Live-cell imaging of HMEC-hTERT1 cells expressing endogenously tagged *RB1-3xFLAG- Clover-sfGFP* and the HDHB CDK sensor ^1^. Cdk-high and Cdk-low cells were grouped as in Fig. 1C. Average traces for the total amount of Rb-3xFLAG-Clover-sfGFP protein (A), nuclear volume (B), the cytoplasmic-to-nuclear intensity ratio of the HDHB sensor (C), and the protein concentration of Rb-3xFLAG-Clover-sfGFP (D, E) are plotted. Traces are aligned by the inflection point of the cytoplasm-to-nuclear ratio of the HDHB sensor, which indicates the initial rise of CDK activity (A-C), or aligned at birth (E). Shaded region denotes the standard deviation. **F**. Representative single cell traces showing the Rb concentration dynamics of Cdk-low and Cdk-high cells. Cells were aligned to the G1/S transition. **G**. Comparing Rb concentrations at birth or at the G1/S transition (Cdk activation point) in Cdk-low and Cdk-high cells.

**Figure S2.**
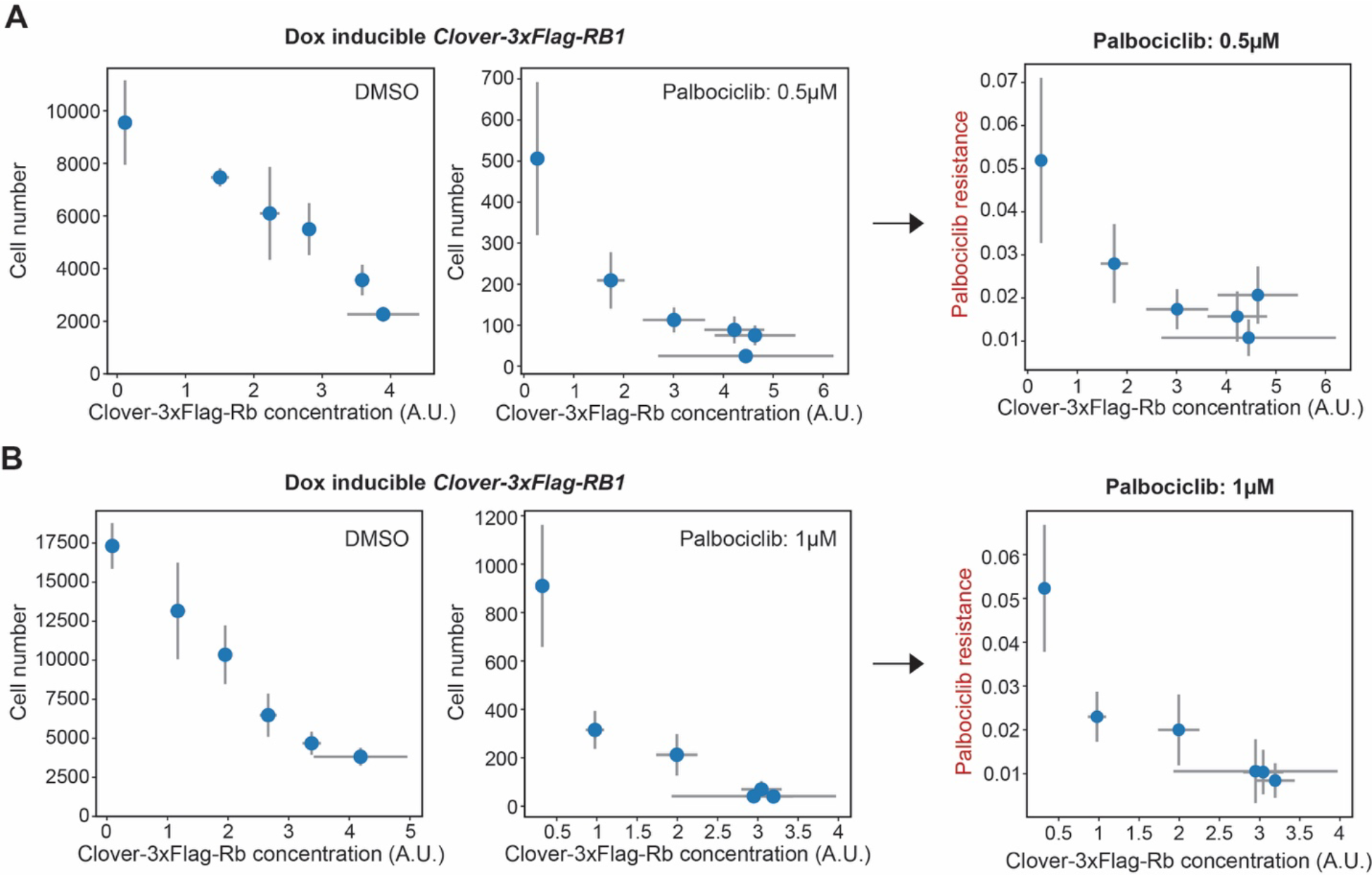
Rb overexpression increases the sensitivity of cells to CDK4/6 inhibition **A-B.** Cell number and normalized cell number of HMEC cells treated with 0.5µM (A) or 1µM (B) Palbociclib for 72 hours. Cells were plated in different doses of doxycycline to induce exogenous Clover-3xFlag-Rb. Drug treatment started the next day and lasted for 72 hours. Then, cells were fixed and the cell number in each well was measured. Normalized cell number is the cell number under Palbociclib treatment divided by the cell number under DMSO treatment. N = 3 biological replicates and the error bars indicate standard deviations.

**Figure S3.**
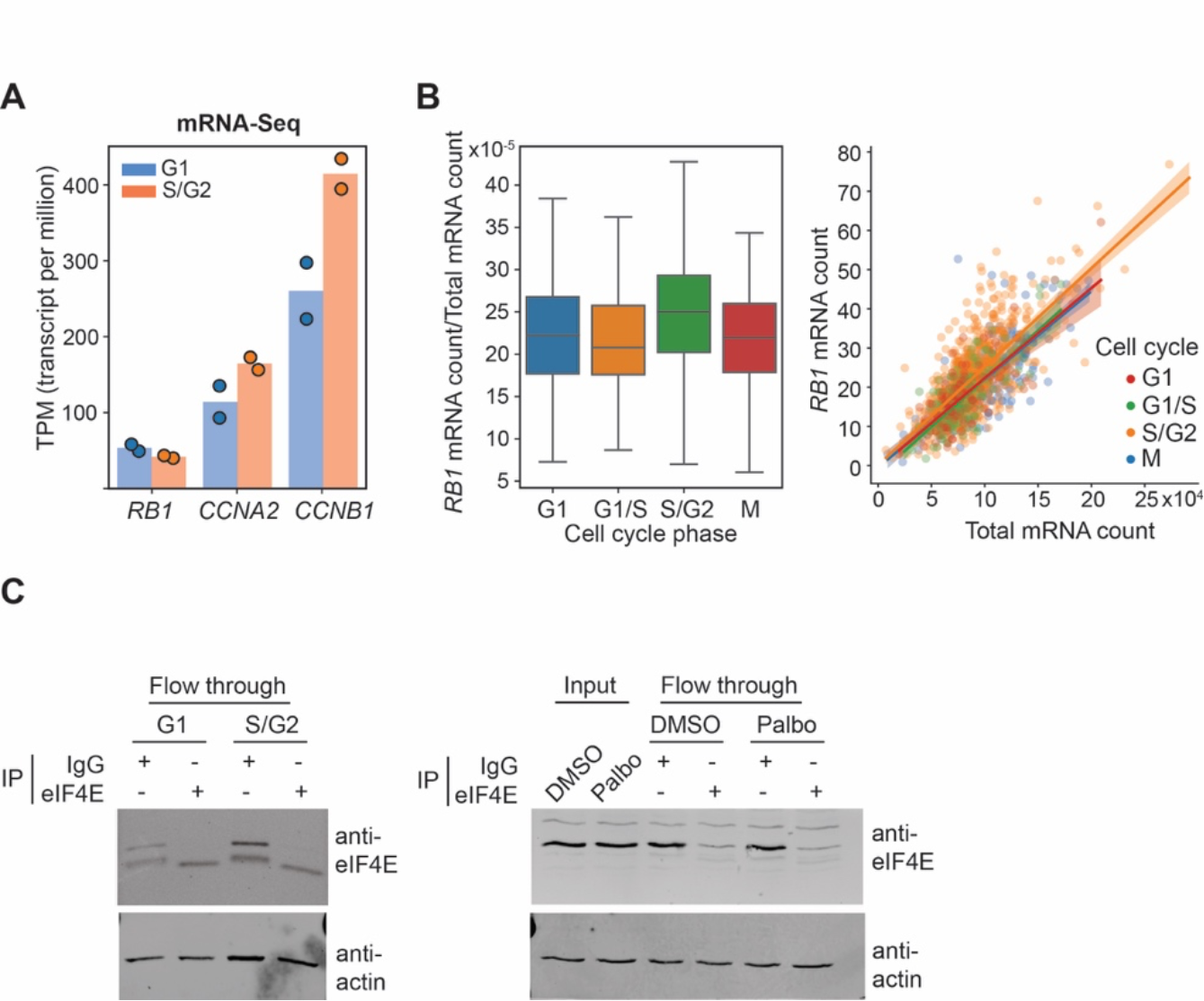
The synthesis rate of *RB1* is relatively constant through the cell cycle. **A.** mRNA-Seq (n = 2) measurements for *RB1* mRNA concentration in G1 and S/G2 HMEC cells sorted using a FUCCI marker. **B.** Analysis of published MERFISH data^35^ for *RB1* mRNA concentrations. (Left panel) the *RB1* mRNA concentration in different cell cycle phases classified using the cell cycle dependent genes identified from the paper. *RB1* mRNA concentration is calculated by dividing the *RB1* mRNA count by the total mRNA count. (Right panel) mRNA count of *RB1* is proportional to the total mRNA count in different cell cycle phases. **C.** Western blot of eIF4E. RIP (RNA binding protein immunoprecipitation) assay for pulling down eIF4E was performed on G1 and S/G2 cell populations or DMSO and Palbociclib (1µM) treated cells.

**Figure S4.**
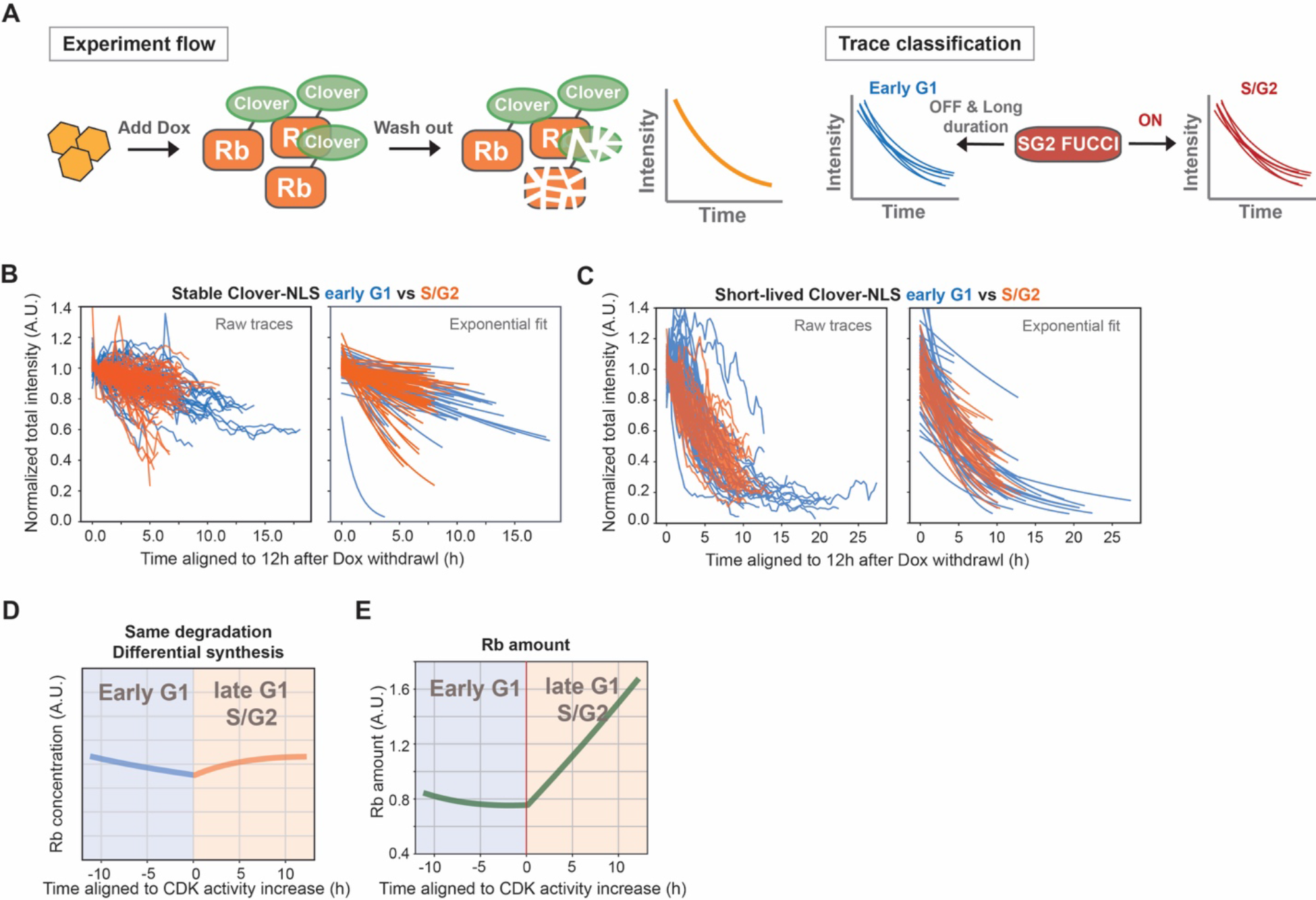
Rb is degraded in early G1 phase. **A.** Schematic of the live-cell imaging approach for measuring the protein degradation rate in different cell cycle phases. **B.** Degradation traces for stable Clover-NLS and the corresponding exponential fits (n=88). **C.** Degradation traces of short-lived Clover-NLS and the corresponding exponential fits (n=57). **D-E**. Mathematical model of Rb concentration and protein amount dynamics during cell cycle progression. (D) Rb concentration dynamics assuming that its degradation rate does not change, but its synthesis rate increases by 20% at the G1/S transition as measured in Figure 2A. (E) Total Rb protein amount dynamics assuming that its degradation rate decreases by 80% at the G1/S transition as measured by live imaging (Figure 2C) and its synthesis rate does not change. Related to Figure 2E.

**Figure S5.**
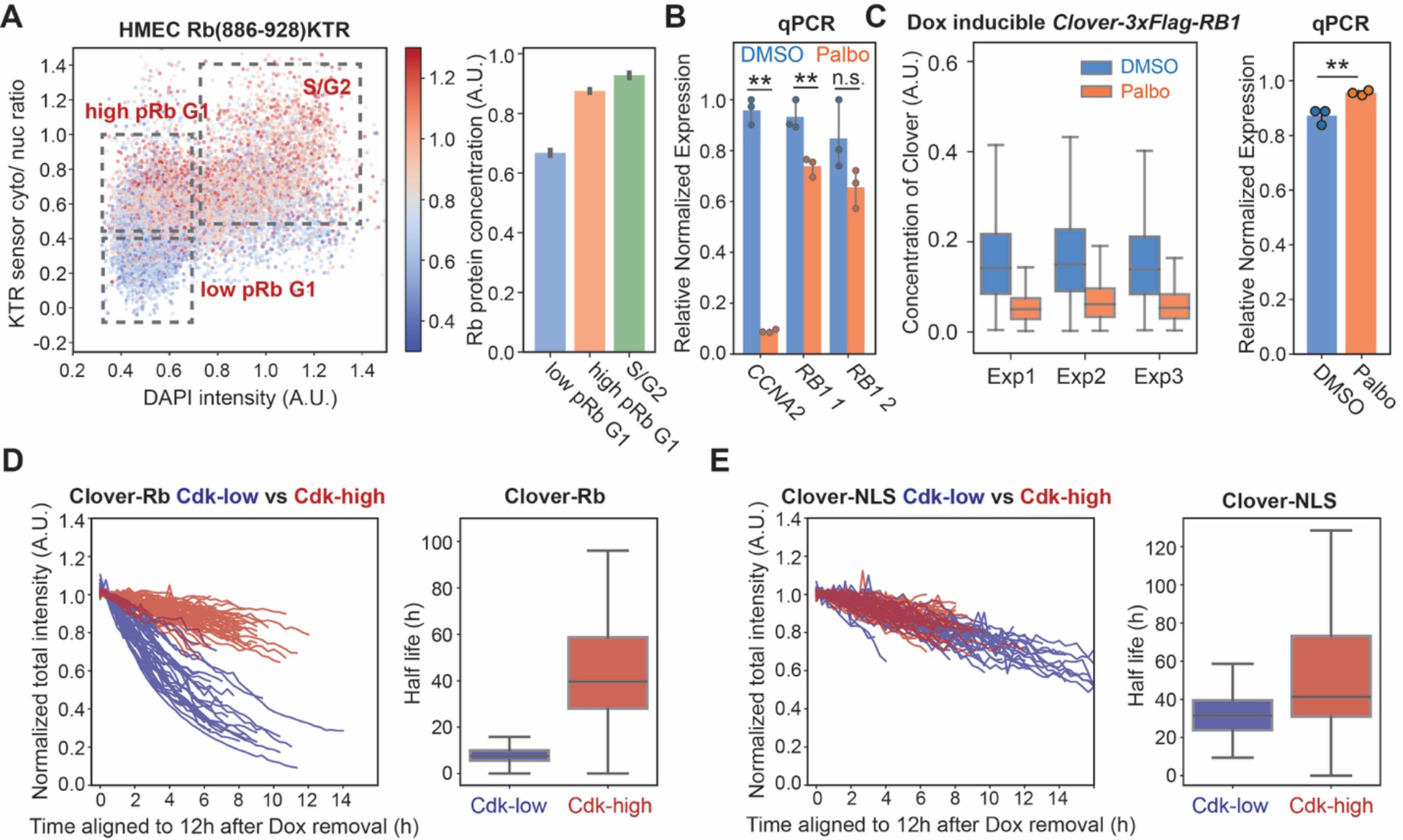
Rb is stabilized by Cdk mediated phosphorylation **A.** Rb concentration in cell populations with different CDK activities. HMEC cells expressing the Rb(886-928) KTR sensor ^36^ were stained with an Rb antibody. (Left panel) KTR sensor cytoplasmic-to-nucleus intensity ratio is plotted against DNA content (DAPI intensity). Dot color indicates Rb concentration. (Right panel) quantification of Rb concentrations in populations with different CDK activities as indicated in regions in the left panel. Bars denote 95% confidence interval for the mean. **B.** *RB1* qPCR measurement of mRNA from cells treated with DMSO or Palbociclib (1µM, 24 hours); related to Fig. 2B. n=3 and dots denote biological replicates. Bars denote standard deviation. **C.** Protein and mRNA concentrations of Clover-3xFlag-Rb. Cells expressing Clover-3xFlag-Rb (Dox induced for 48 hours) were treated with DMSO or Palbocliclib (1µM) for 24 hours. The protein concentration of Clover-3xFlag-Rb (left panel) was measured by flow cytometry. The concentration was defined as Clover fluorescent intensity divided by side scatter area (SSC-A). The mRNA concentration of Clover-3xFlag-Rb (right panel) was measured using qPCR with primers targeting Clover. **D-E.** (Left panel) The degradation traces of Clover-3xFlag-Rb protein (D) or Clover-NLS (E) after Dox withdrawal. The traces were classified into Cdk-low and Cdk-high cells based on the HDHB Cdk sensor. (Right panel) Distribution of half-lives estimated from the exponential fit. Box plot indicates 5th, 25th, median, 75th and 95th percentiles.

**Figure S6.**
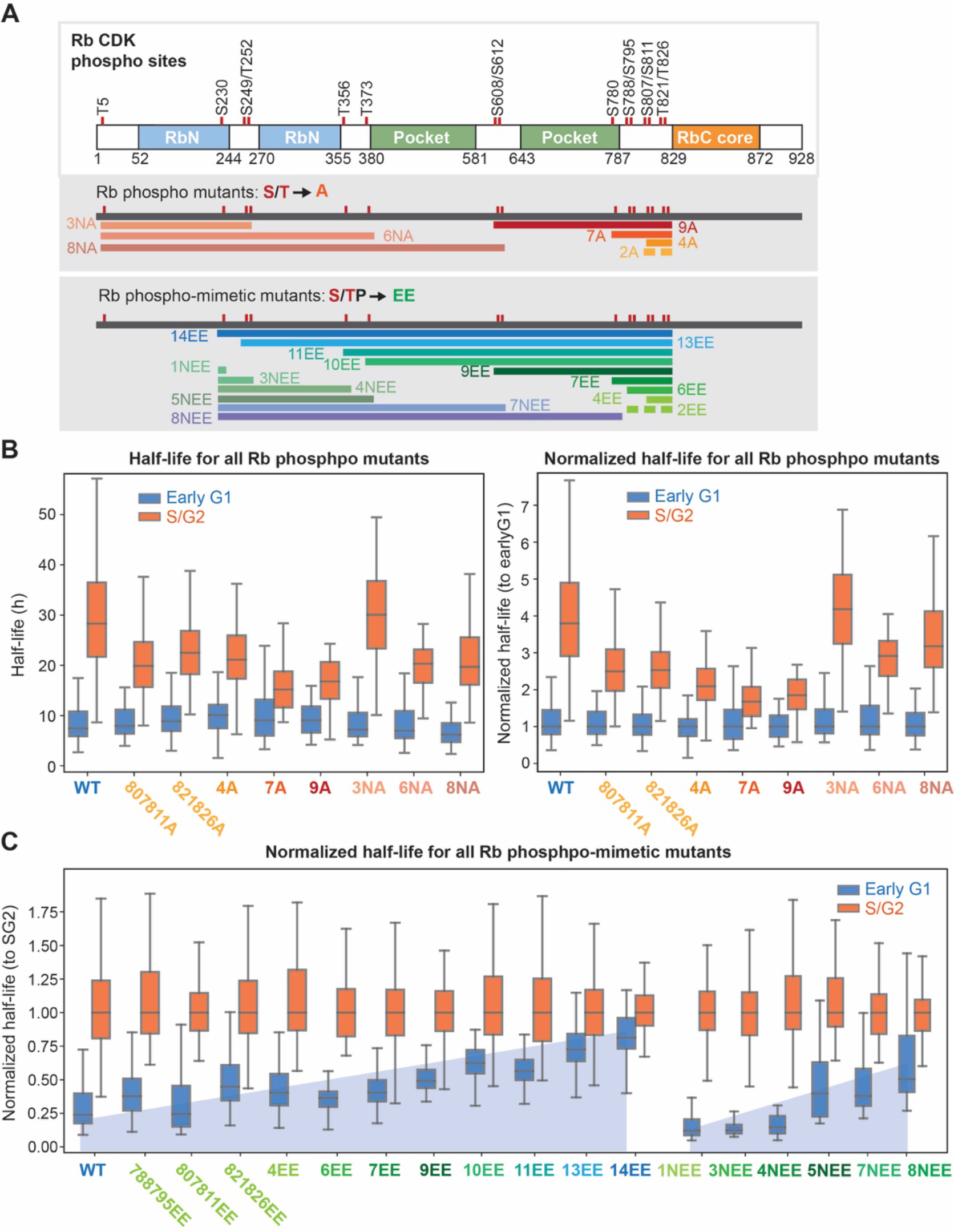
Half-life measurements of Rb phospho-mutants and phosphomimetic mutants A. Schematics of the CDK sites on Rb that were mutated in the Rb phospho-mutants and the Rb phospho-mimetic mutants used in this study. **B.** The half-lives (left) and normalized half-lives (right) for all the Rb phospho-mutants measured using the live-cell imaging approach. The normalized half-lives were calculated by normalizing to the half-life in early G1 for each Rb protein variant. **C.** Normalized half-lives of all the Rb phospho-mimetic mutants measured using the live-cell imaging approach. Related to Figure 3E, F. Here, the normalized half-lives were calculated by normalizing to the half-life in S/G2 for each Rb protein variant.

**Figure S7.**
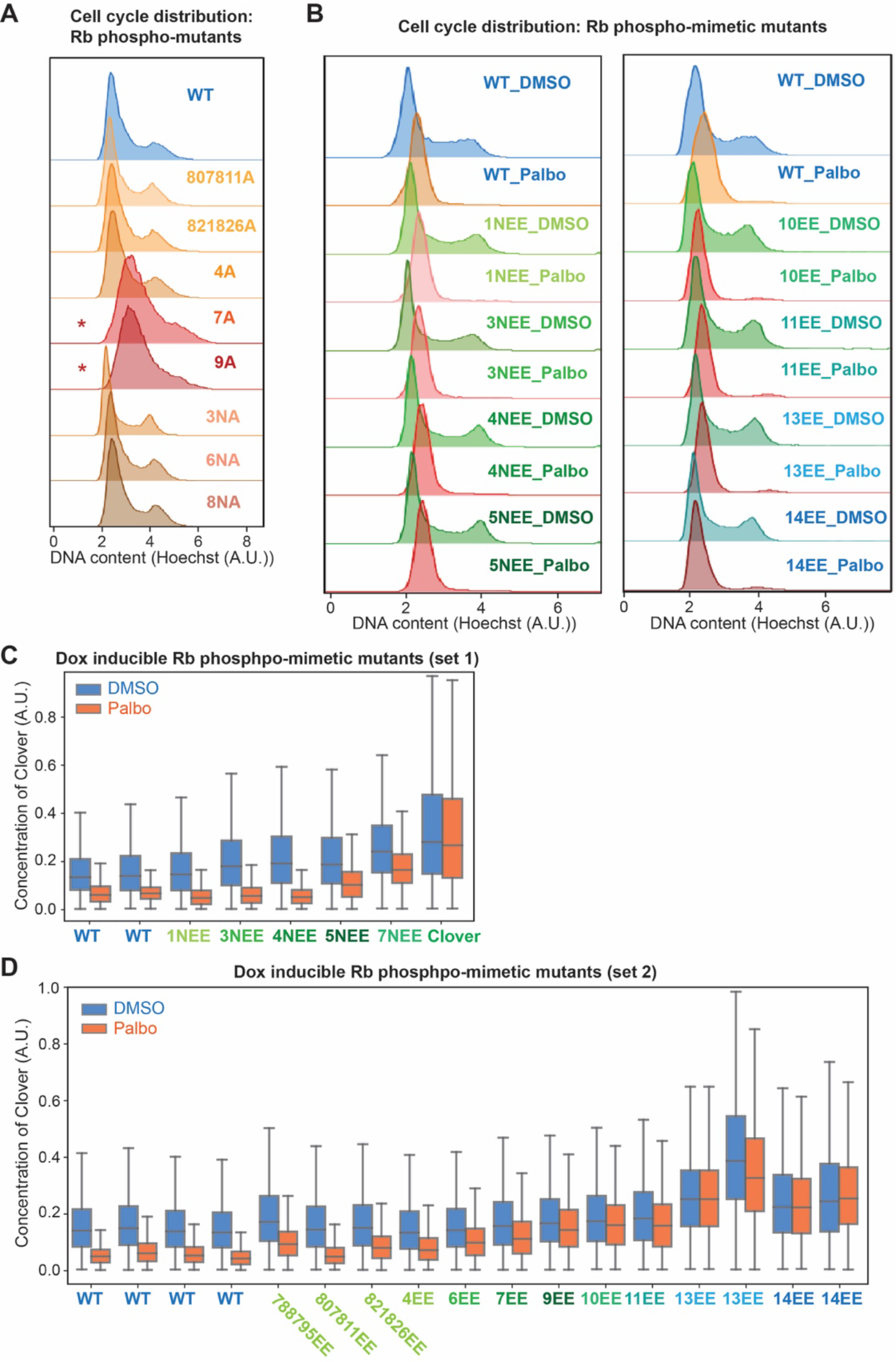
Cdk phospho-site mutations affect Rb stability A. Cell cycle distribution for cells expressing different Rb phospho-mutants. Cells expressing Clover-3xFlag-Rb phospho-mutants (Dox induced for 48 hours) were incubated with Hoechst for 30 minutes to stain DNA, and then analyzed by flow cytometry. Red stars indicate that cells expressing Rb7A and Rb9A are arrested in G1. **B.** Cell cycle distribution for cells expressing Rb phospho-mimetic mutants. Cells expressing Clover-3xFlag-Rb phospho-mimetic mutants (Dox induced for 48 hours) were treated with DMSO or Palbocliclib (1µM) for 24 hours. Then the cells were incubated with Hoechst for 30 minutes to stain DNA, and analyzed by flow cytometry. **C-D**. The concentrations of Clover-3xFlag-Rb phospho-mimetic mutants (induced by Dox for 48 hours) after 24 hours of DMSO or Palbociclib (1µM) treatment, as measured by flow cytometry. (**C**) shows the mutations made from the N-terminus of Rb, and (**D**) shows the mutations made from the C-terminus of Rb.

**Figure S8.**
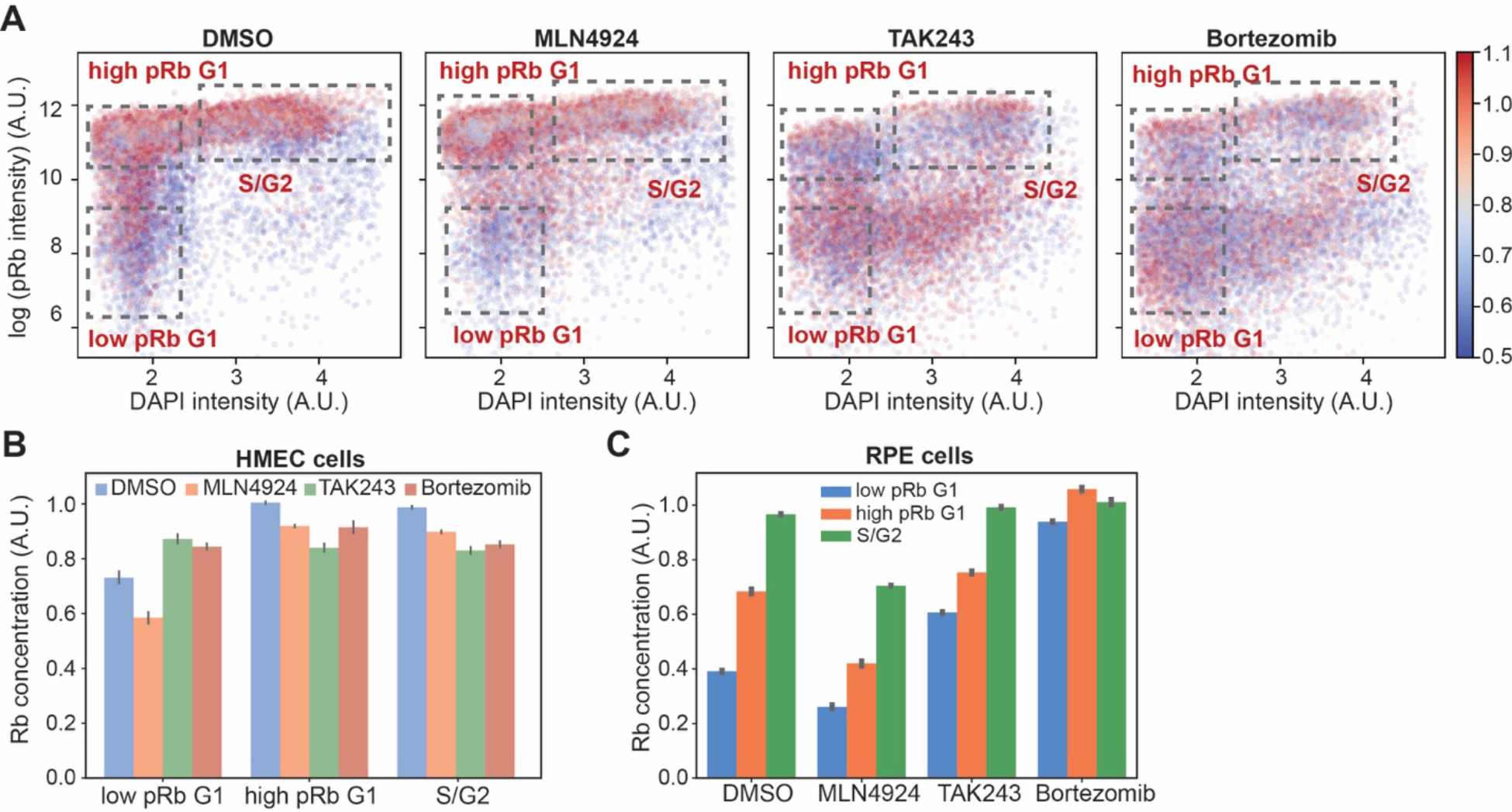
Un/hypo-phosphorylated Rb is degraded through the Ubiquitin-proteasome system **A.** HMEC cells were treated with the indicated drugs for 5 hours, fixed, and stained with phospho-Rb (S807/811) and Rb antibodies. The Rb concentration is calculated by dividing total Rb intensity by nuclear area^3/2^. The dot color indicates Rb concentration. **B**. Rb concentration in different phospho-Rb populations of HMEC cells shown in A. **C.** Rb concentration in different phospho-Rb populations of RPE-1 cells after treatments with the indicated drugs for 6 hours, as measured by flow cytometry. Rb concentration was calculated by dividing the Rb total intensity with the side scatter area (SSC-A).

**Figure S9.**
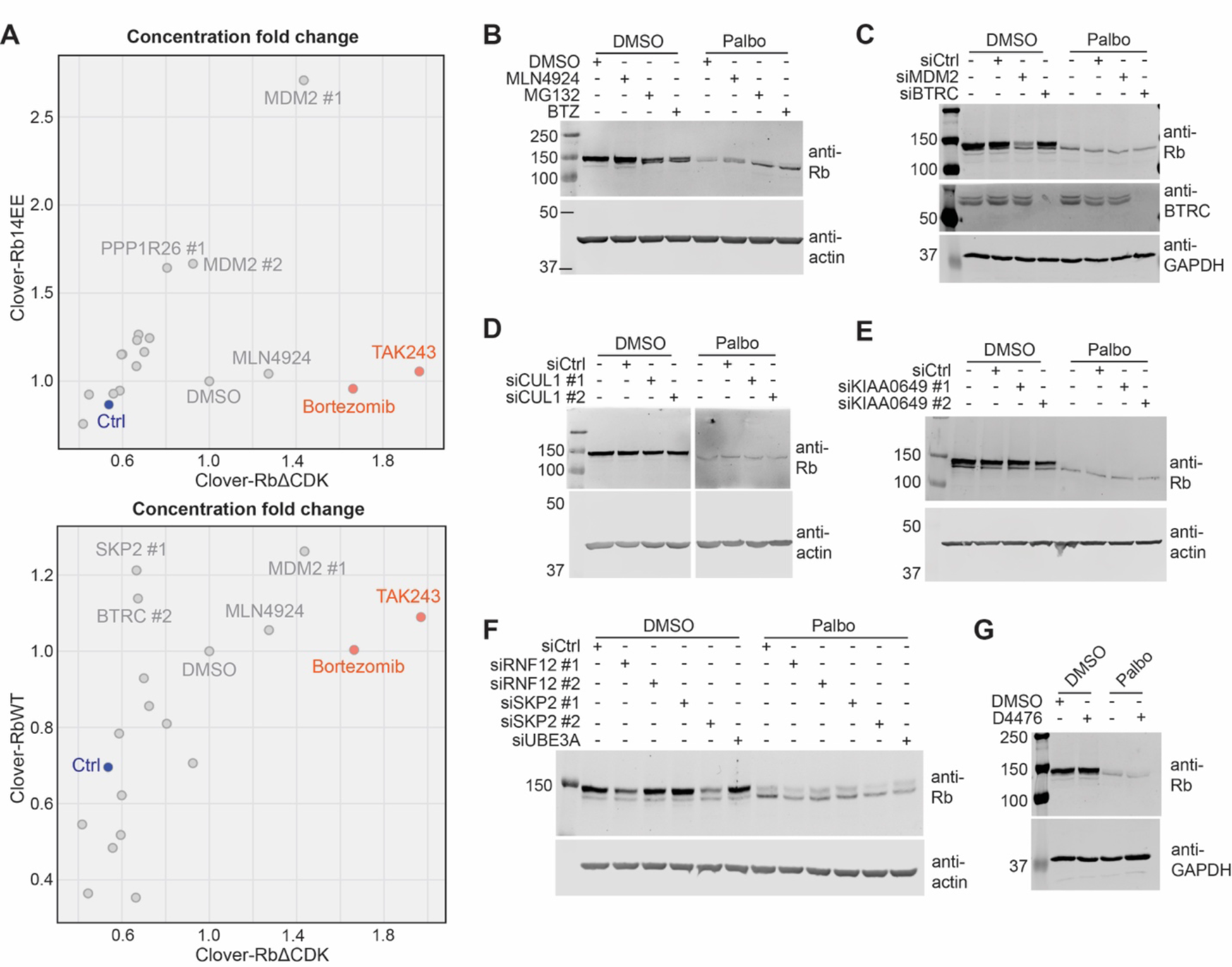
Examination of E3 ligases reported to target Rb **A.** Comparison of concentration fold change after siRNA treatment. HMEC cells expressing Clover-3xFlag-RbΔCDK, Clover-3xFlag-RbWT, or Clover-3xFlag-Rb14EE (Dox induced for 48 hours) were transfected with siRNAs against the published E3 ligase genes for 48 hours. Then, the cells were fixed and imaged. The concentration fold change is calculated by dividing the Clover concentration of the treatment well by the concentration in the DMSO treated well. Red dots indicate the positive controls, *i.e.*, TAK243 and Bortezomib treatments (5 hours of treatment), and the blue dot indicates the siCtrl treated sample. **B-F**. The published E3 ligase genes for Rb were also examined by western blot analysis. HMEC cells were transfected with the corresponding siRNAs for 24 hours, and then treated with DMSO or Palbociclib (1µM) for 24 hours. Cells were then harvested for western blotting with Rb antibodies. The lower band in the Rb western blots represents un/hypo-phosphorylated Rb. **G.** The effect of CK1 on Rb half-life was examined using western blot analysis. Cells were treated with DMSO vs Palbociclib (1µM) and DMSO vs D4476 (25µM CK1 inhibitor), and then the cells were lysed for western blotting.

**Figure S10.**
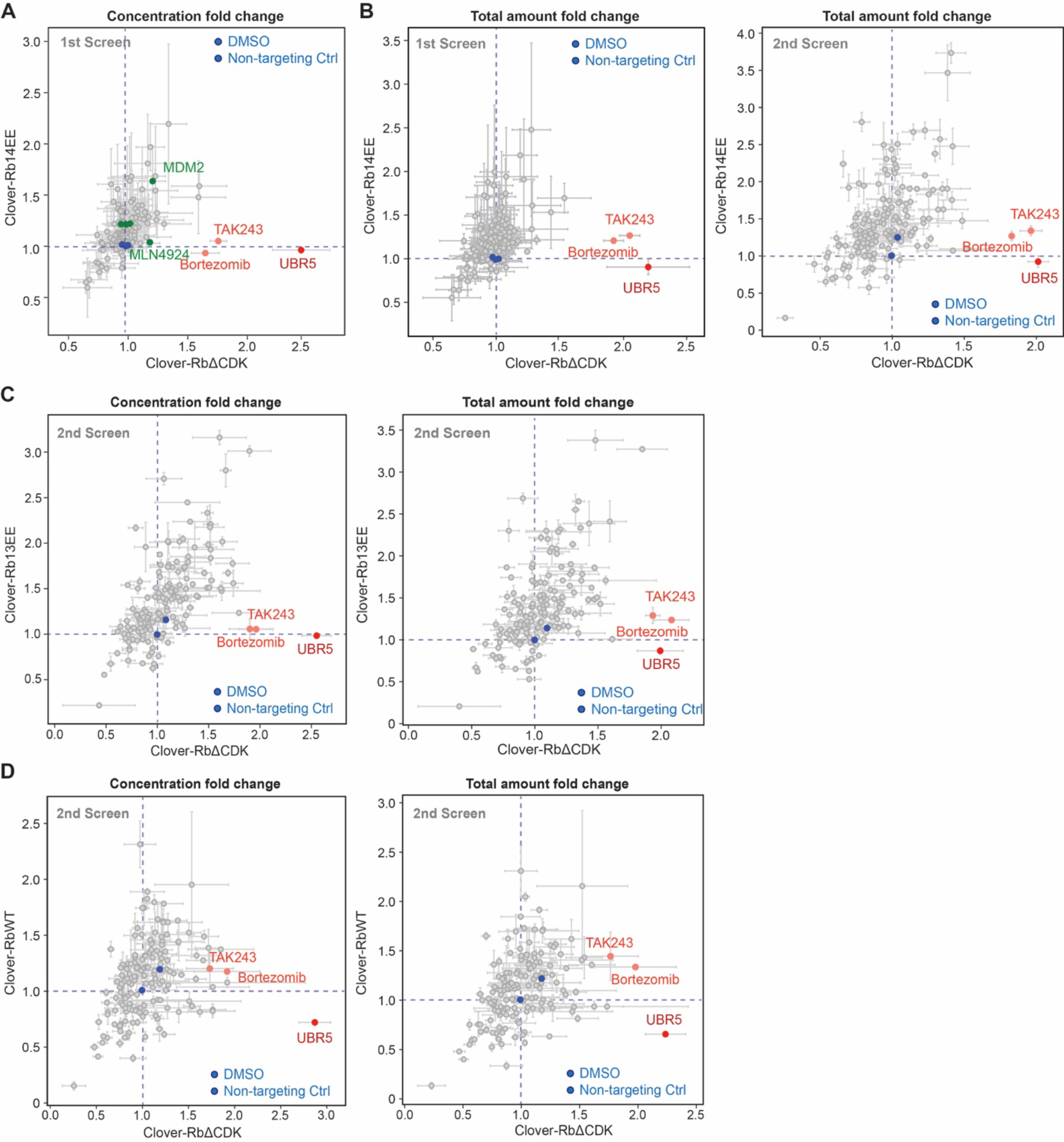
UBR5 is identified as a candidate E3 ligase targeting Rb in siRNA screens **A.** Same plot as Figure 4D (1st siRNA screen), except that the green dots are added to indicate the previously reported E3 genes for Rb. **B.** The total amount fold changes of Clover-3xFlag- RbΔCDK and Clover-3xFlag-Rb14EE are plotted. Related to Figure 4D, E. **C.** Concentration fold changes and total amount fold changes of Clover-3xFlag-RbΔCDK and Clover-3xFlag-Rb13EE from the 2nd siRNA screen. n = 2 biological replicates. **D.** Concentration fold changes and total amount fold changes of Clover-3xFlag-RbΔCDK and Clover-3xFlag-RbWT from the 2nd siRNA screen. n = 2 biological replicates.

**Figure S11.**
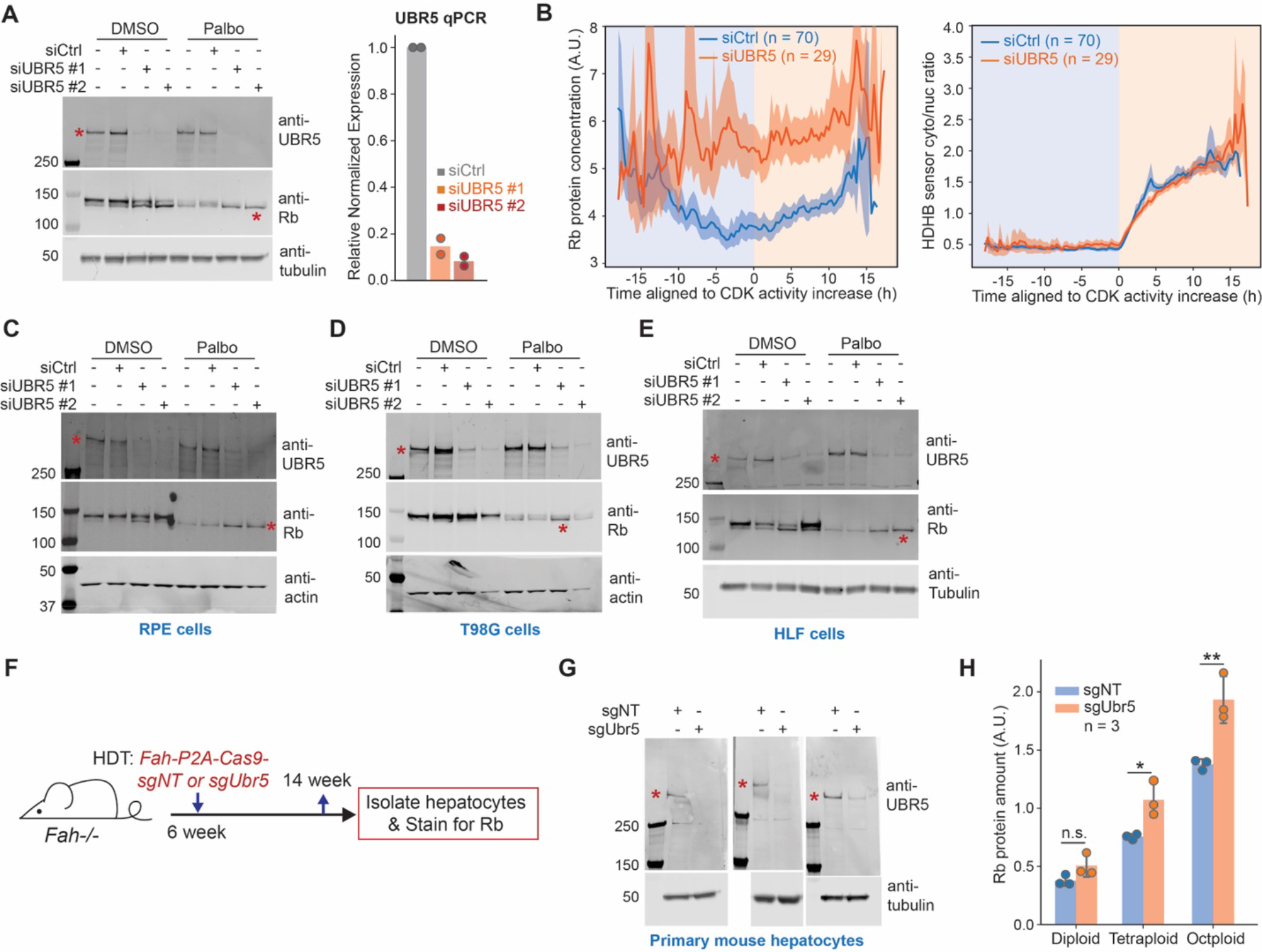
UBR5 is responsible for Rb degradation in multiple cell types A. Left panel: Immunoblot of Rb after UBR5 knockdown by siRNA. Cells were treated with siRNAs for 24 hours, and then treated with DMSO or Palbociclib (1µM) for 24 hours before harvest. The lower band in the Rb blot indicates the un/hypo-phosphorylated Rb (marked by red star). Right panel: qPCR validation of UBR5 knockdown efficiency. HMEC cells were transfected with siRNA for 48 hours. **B.** Live imaging validation of Rb concentration dynamics during cell cycle progression. HMEC cells expressing an endogenously tagged *RB1-3xFLAG- Clover-sfGFP* and an HDHB CDK sensor ^1^ were transfected with UBR5 siRNA for 24 hours and then imaged. Average traces for Rb-3xFLAG-Clover-sfGFP concentration (left panel) or the cytoplasmic-to-nuclear intensity ratio of the HDHB sensor (right panel) are plotted; solid lines indicate mean values for each time point and shaded regions indicate the standard deviation of the mean. Traces are aligned using the HDHB sensor to identify the time that CDK activity begins to increase. **C-E.** Western blot validation of the role of UBR5 in Rb degradation in multiple cell lines (**C.** RPE cells, **D.** T98G cells, **E.** HLF cells). Cells were transfected with siRNAs against UBR5 or Ctrl siRNA. 24 hours later, DMSO or Palbociclib (1µM) were added for another 24 hours. Then, cells were lysed for western blot analysis. The lower band of the Rb blot is marked by a red star, which represents the un/hypo-phosphorylated Rb. **F**. Experimental flow of the *Fah-/-* mouse liver model. 6-week old *Fah-/-* mice were hydrodynamically transfected with plasmids carrying an *Fah-P2A-Cas9-sgNT* transposon or an *Fah-P2A-Cas9-sgUbr5* transposon, together with a transposase plasmid. 8 weeks later, hepatocytes were isolated from the mice for downstream analysis. **G.** Western blot validation of UBR5 knockout efficiency in primary hepatocytes. *Fah-/-* mice were injected with transposons carrying *Cas9-sgNT* or *Cas9- sgUbr5*. Primary hepatocytes were isolated from the mice, plated, and then lysed for western blot analysis. n = 3 pairs of mice. **H.** Total Rb amount in the low-phospho-Rb population of the primary hepatocytes isolated from mice receiving *Fah-P2A-Cas9-sgNT* or *Fah-P2A-Cas9- sgUbr5* transposons. n = 3 pairs of mice. Error bar indicates standard deviation. Related to Figure 5D.

**Figure S12.**
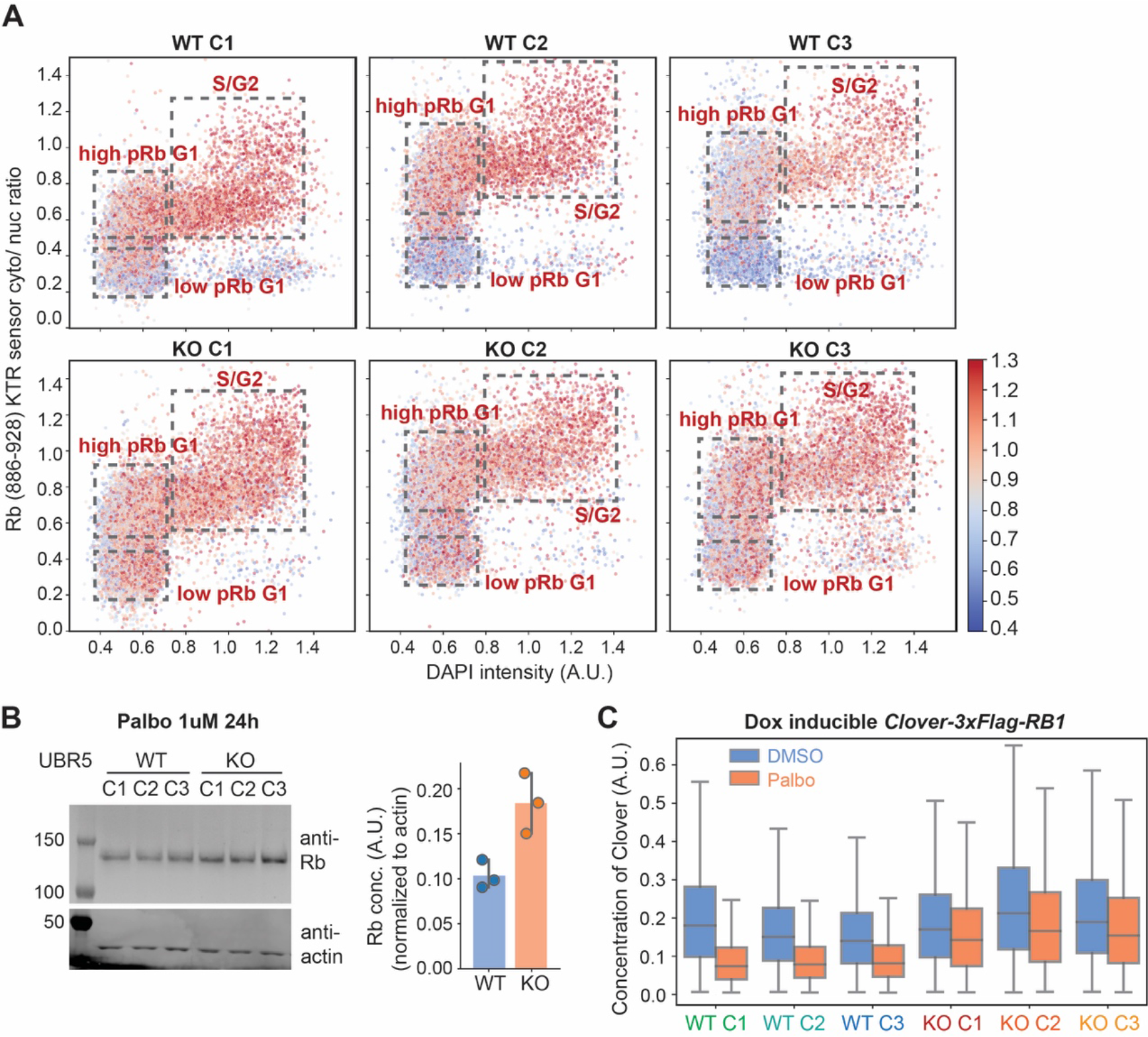
Rb concentration is increased in the low-phospho-Rb populations of *UBR5 KO* clonal cell lines **A**. Rb staining of *UBR5 WT* or *KO* clonal cell lines expressing the Rb (886-928) KTR sensor ^36^. The cytoplasmic-to-nucleus intensity ratio of the KTR sensor is plotted against DNA content (DAPI). The dot color indicates the Rb concentration. Related to Fig. 4B. **B.** Western blot analysis of Rb concentration after Palbociclib treatment in UBR5 clonal cell lines. 6 UBR5 clonal cell lines (3 WT, 3 KO) were treated with DMSO or Palbociclib (1µM) for 24 hours, and then lysed for western blotting. Quantification of Rb concentration (Rb band intensity normalized to the actin band intensity) is shown on the right. Dots denote individual clones and error bars denote the standard deviation. **C.** Flow cytometry analysis of Clover-3xFlag-Rb concentration after Palbociclib treatment. 6 UBR5 clonal cell lines (3 WT, 3 KO) expressing Clover-3xFlag- RbWT (Dox induced for 48 hours) were treated with DMSO or Palbociclib (1µM) for 24 hours, and then cells were analyzed using flow cytometry. The concentration of Clover-3xFlag-RbWT was defined as Clover fluorescent intensity divided by side scatter area (SSC-A).

**Figure S13.**
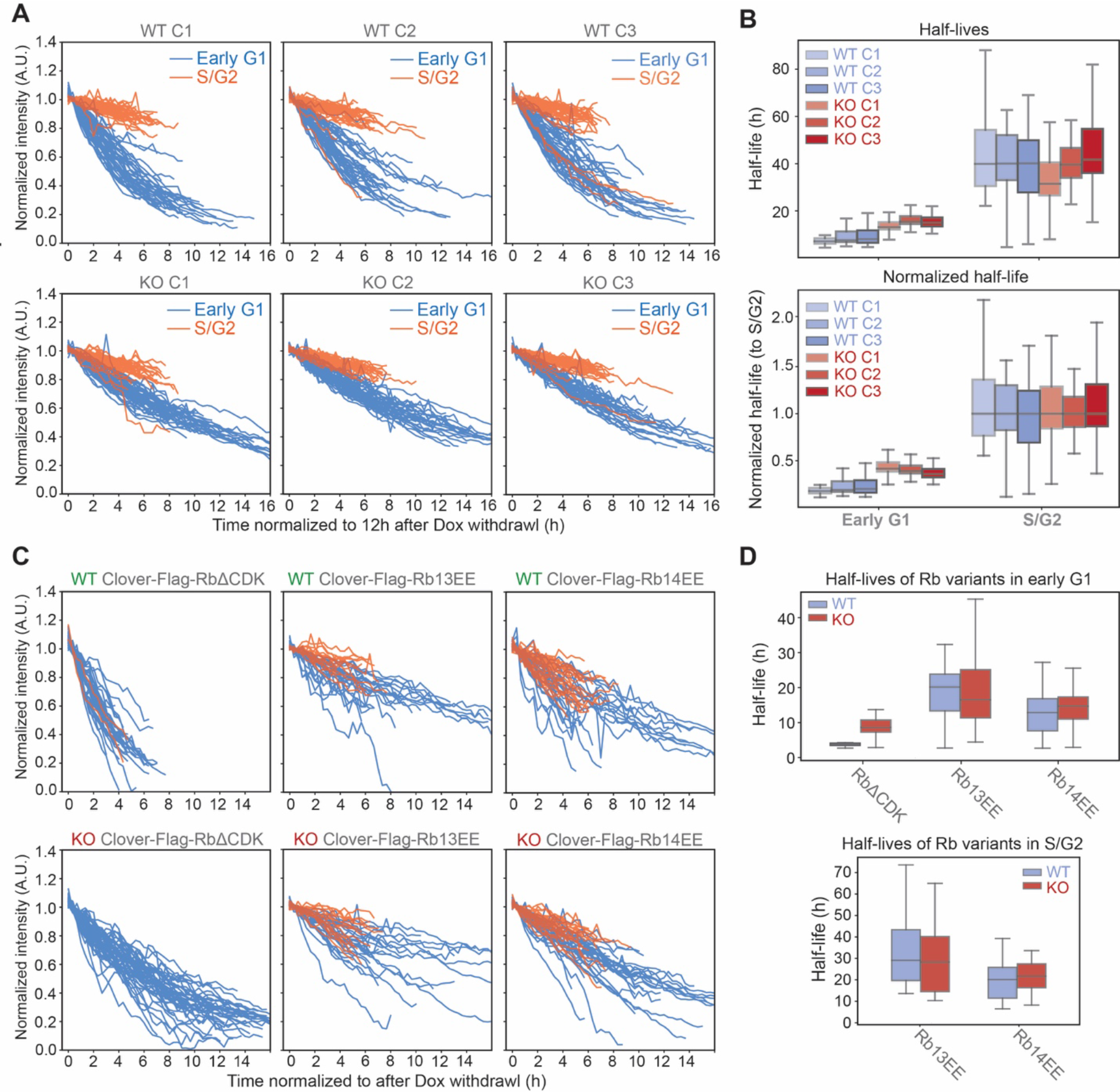
Un-phosphorylated Rb is stabilized in *UBR5 KO* cells A. The degradation traces of Clover-3xFlag-RbWT protein after Dox withdrawal in 6 UBR5 clonal cell lines (3 WT, 3 KO). The traces were classified into early G1 phase or S/G2 phase based on the FUCCI cell cycle reporter. **B.** The half-life distributions (upper panel) and normalized half-lives (lower panel, normalized to the median of S/G2 half-life) calculated from the degradation traces shown in (**A**). **C**. The degradation traces of Clover-3xFlag-RbΔCDK, Clover-3xFlag-Rb13EE, and Clover-3xFlag-Rb14EE protein after Dox withdrawal in UBR5 WT and KO cell lines. The traces were classified into early G1 phase or S/G2 phase based on a FUCCI cell cycle reporter. **D**. The half-life distributions of the Rb variants in (C) in early G1 phase (upper panel) or in S/G phase (lower panel) calculated from the degradation traces shown in (C).

**Figure S14.**
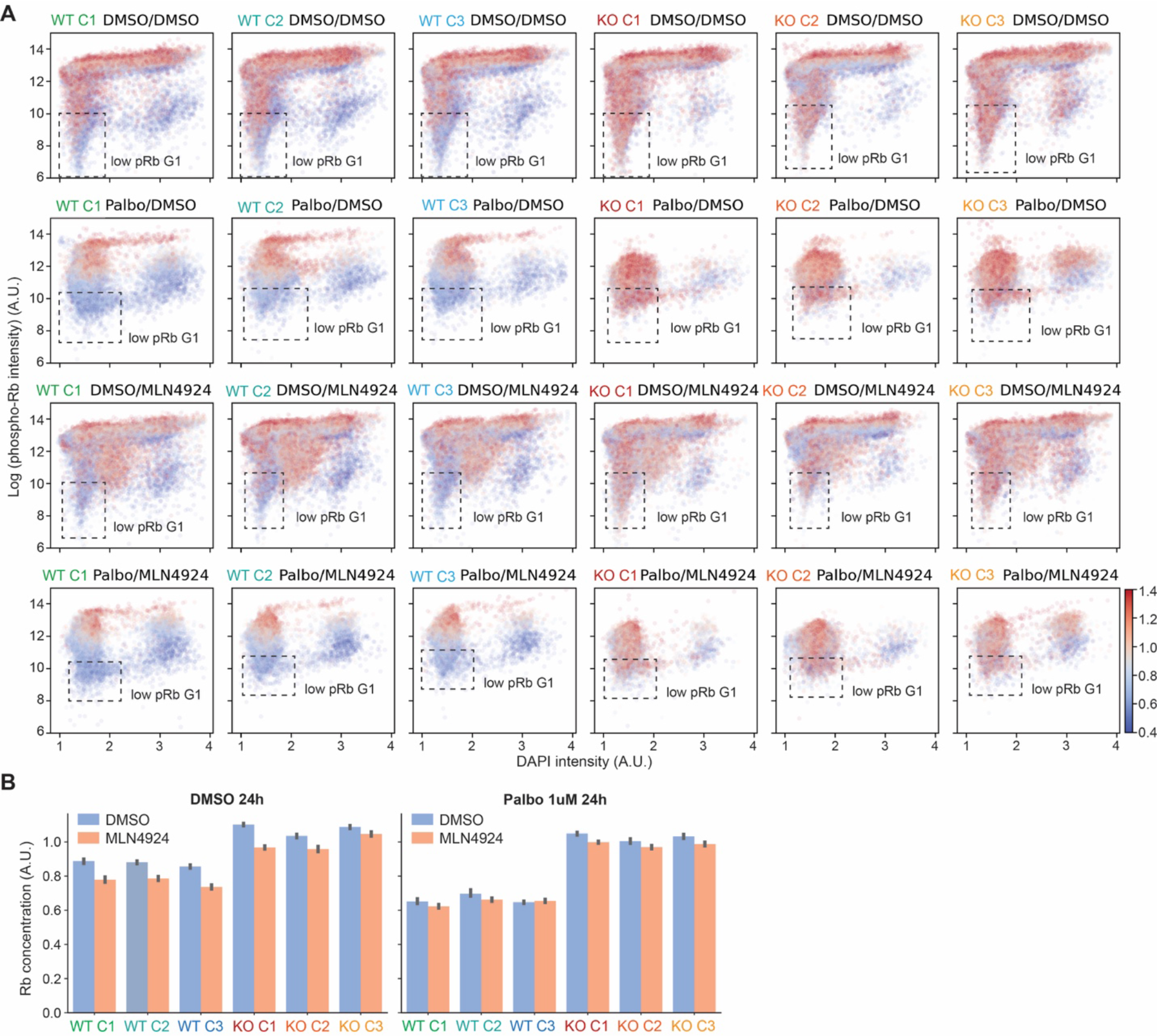
The Rb concentration is increased in the low-phospho-Rb populations of *UBR5 KO* clonal cell lines. **A.** phospho-Rb concentration is plotted against DNA content in UBR5 clonal cell lines. 6 UBR5 clonal cell lines (3 WT, 3 KO) were treated with DMSO or Palbociclib (1µM) for 24 hours, and then the cells were treated with DMSO or MLN4924 (1µM) for 5 hours before fixation. Cells were stained with phospho-Rb (S807/811) and Rb antibodies, followed by imaging. The dot color indicates the Rb concentration. The dashed line box indicates the low pRb G1 population that has low concentration of phospho-Rb. Plot titles indicate the cell genotype and the treatments administered. *E.g.*, Palbo/DMSO indicates that 1 µM Palbociclib was administered before DMSO. **B.** Quantification of the Rb concentration in the low pRb G1 populations of cells in (A). Error bar indicates the 95% confidence interval of the mean.

**Figure S15.**
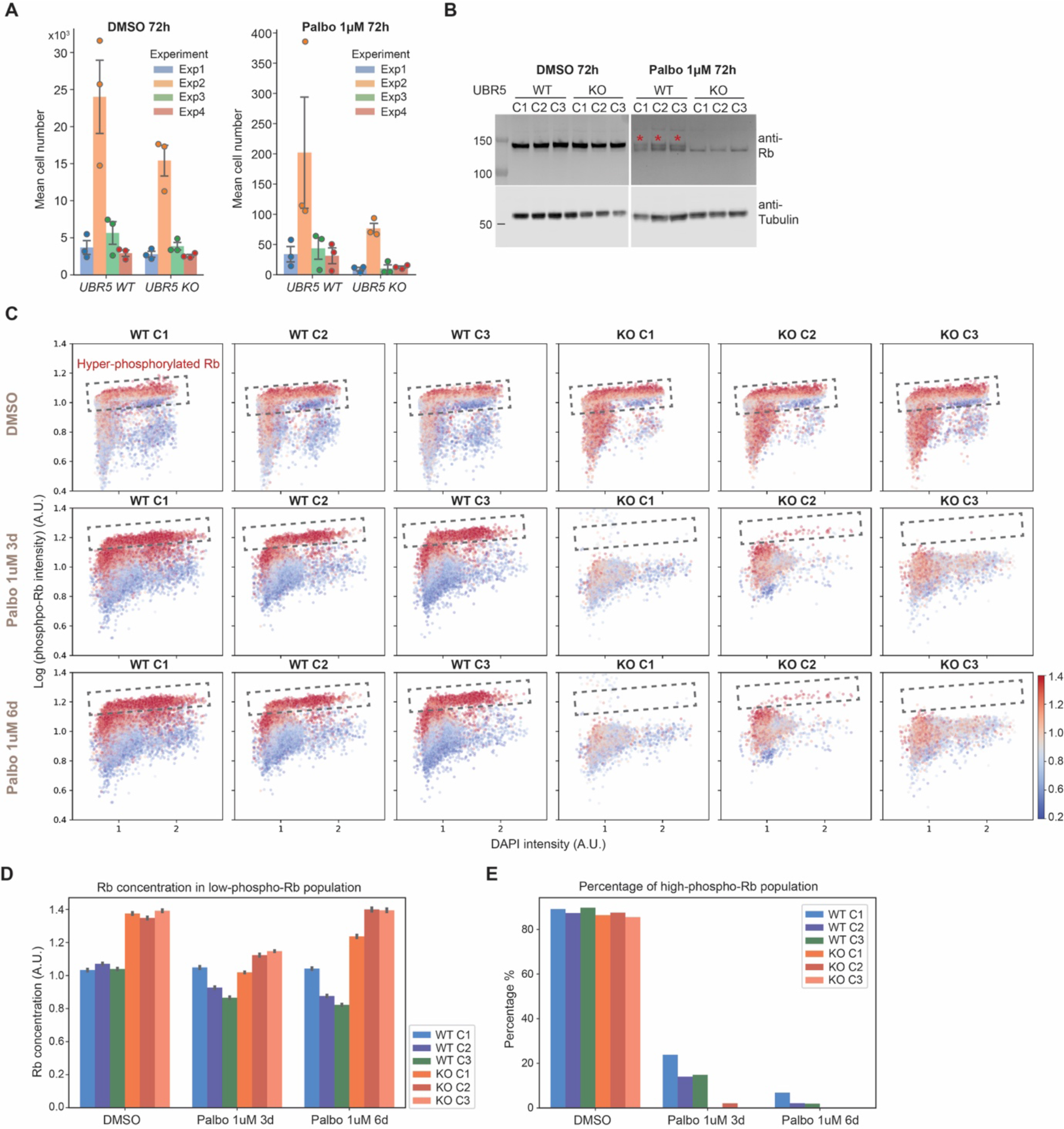
***UBR5 KO* cells are sensitized to Palbociclib treatment A.** The absolute cell number counts for 6 UBR5 clonal cell lines (3 WT, 3 KO) after DMSO or Palbociclib (1µM) treatment for 72 hours; related to Fig. 4E. **B-C.** Western blot (**B**) and immunostaining (**C**) analysis of phospho-Rb (S807/811) in 6 UBR5 clonal cell lines (3 WT, 3 KO) after DMSO or Palbociclib (1µM) treatment for 3 days or 6 days. The hyper-phosphorylated Rb population is marked using red stars (the upper band in (B)) or dashed lined boxes (in (C)). **D**. Quantification of the Rb concentration in the low pRb G1 populations of cells in (C). Error bar indicates the 95% confidence interval of the mean. **E**. Quantification of the percentage of hyper- phosphorylated Rb population of cells in (C).

**Figure S16.**
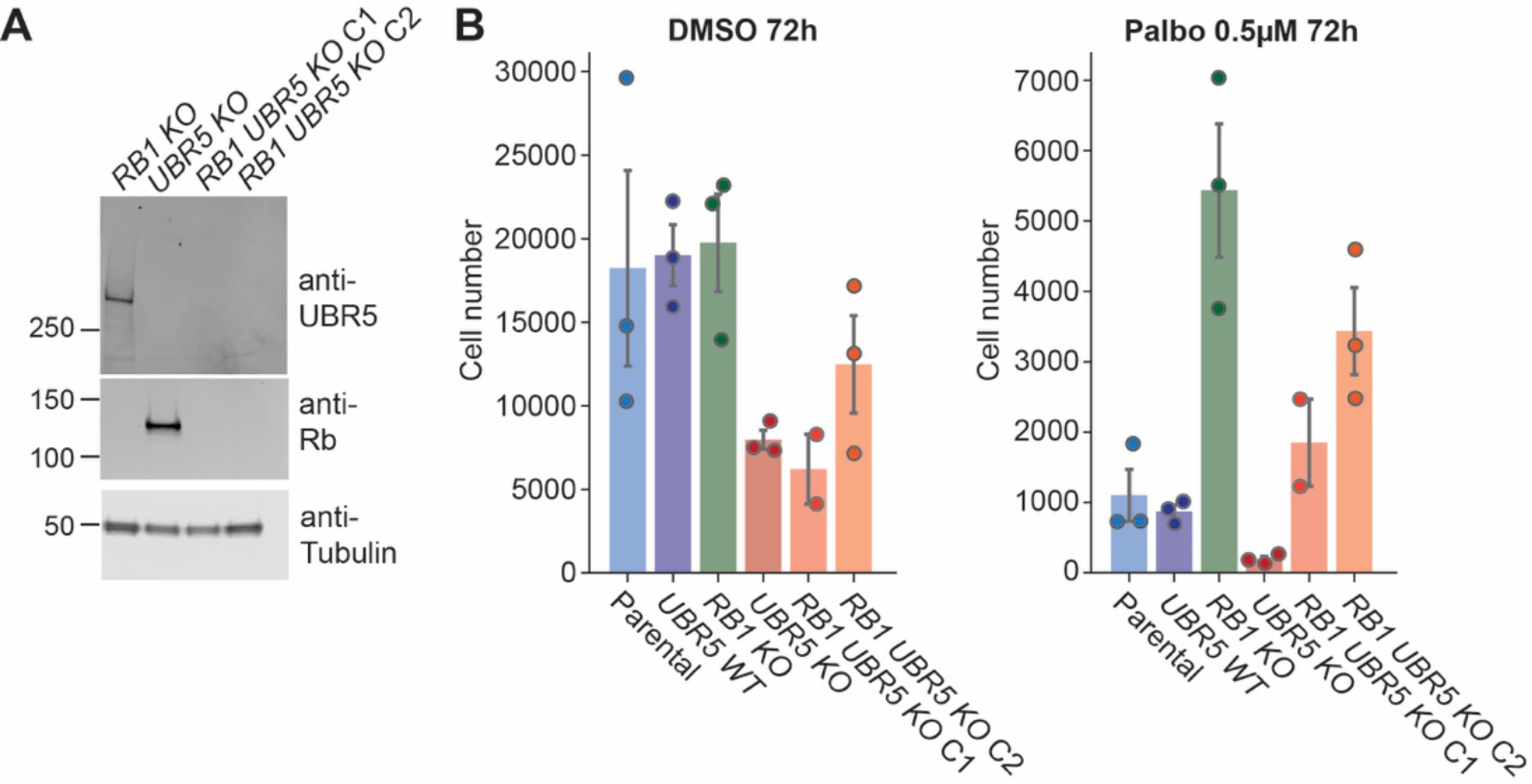
The increased sensitivity to Palbocilib treatment in *UBR5 KO* cells depends on Rb. **A.** Western blot validation of UBR5 and Rb double KO cell lines. **B**. The absolute cell number count for wild-type, *UBR5 KO*, *RB1 KO*, and *UBR5 RB1 KO* cells after DMSO or Palbociclib (0.5µM) treatment for 72 hours; related to Fig. 4F. Bars denote the standard deviation of the mean.

**Figure S17.**
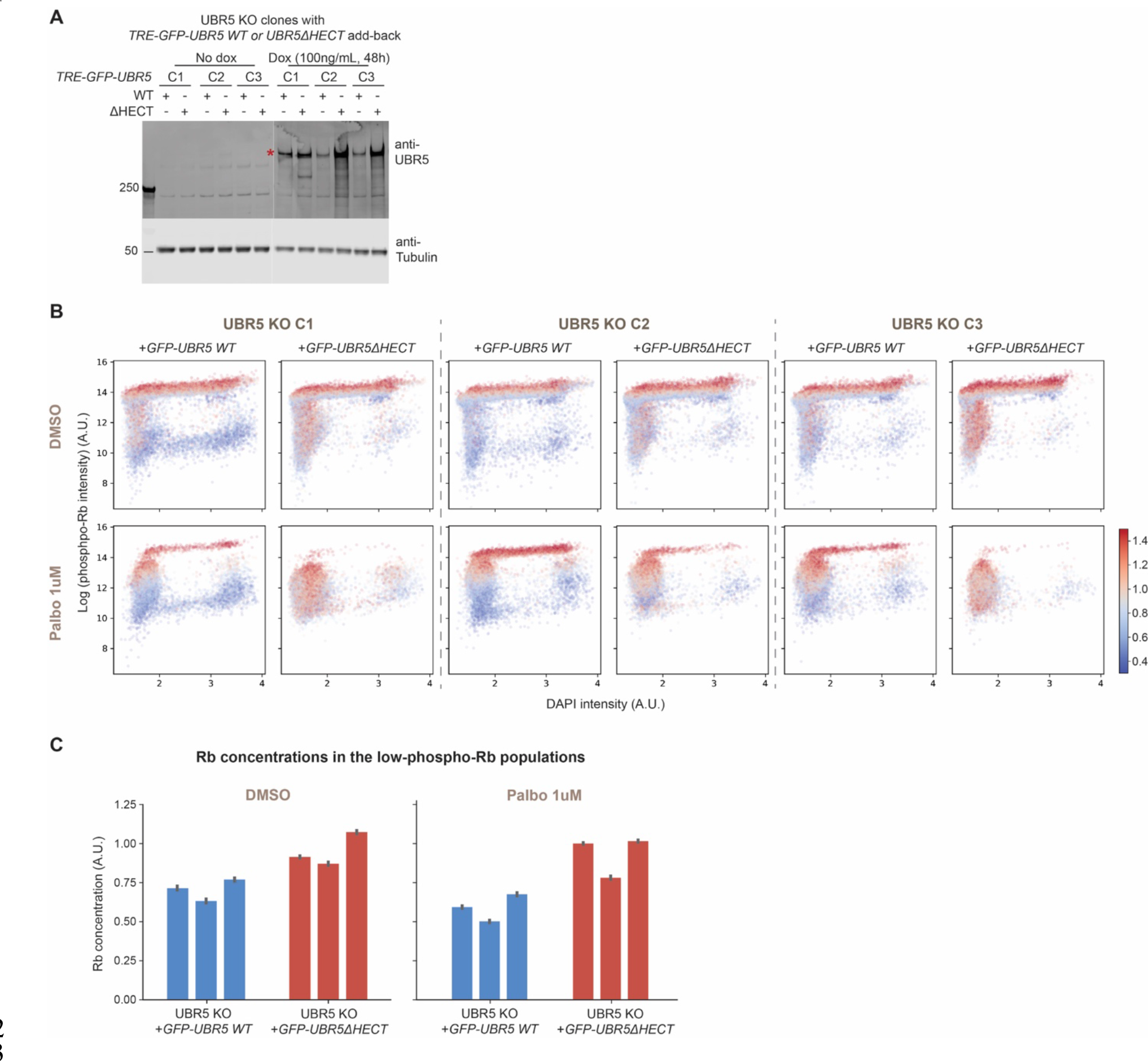
Adding back WT UBR5, but not mutant UBR5, decreases Rb concentration in *UBR5 KO* cells A. Western blot validation of *UBR5 KO* cell lines (3 KO clones) with UBR5WT or UBR5ΔHECT add-back using the Dox inducible system. Cells were induced with Dox (100ng/ml) for 48 hours and then lysed for western blot analysis. **B**. immunostaining analysis of phospho-Rb (S807/811) in 3 *UBR5 KO* clonal cell lines with UBR5WT or UBR5ΔHECT add-back (as described in A) after DMSO or Palbociclib (1µM) treatment for 24 hours. The dot color indicates the Rb concentration. **C**. Quantification of the Rb concentration in the low-pRb G1 populations of cells in (B). Error bar indicates the 95% confidence interval of the mean.

**Figure S18.**
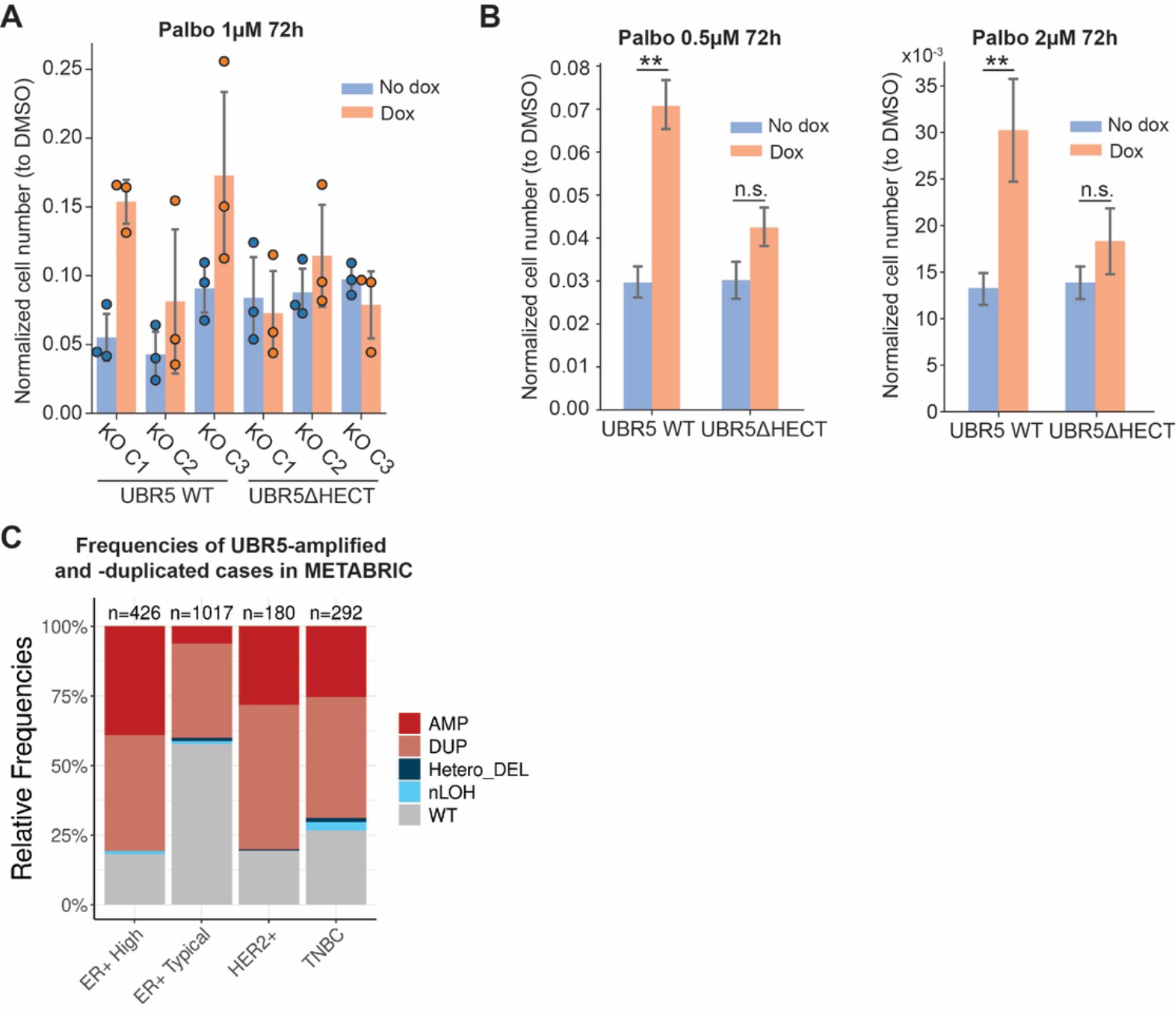
Adding back WT URB5 rescues the Palbociclib-sensitivity phenotype. **A.** Normalized cell number of *UBR5 KO* cells with UBR5 WT or UBR5ΔHECT addback after Palbociclib (0.5µM) treatment for 72 hours (related to Fig. 4G). **B.** Normalized cell number of *UBR5 KO* cells with UBR5 WT or UBR5ΔHECT addback after Palbociclib (0.5µM or 2µM) treatment for 72 hours. The results are the average of 3 *UBR5 KO* clonal cell lines with different UBR5 addback and 3 biological replicates. All error bars denote standard deviations. **C.** Frequencies of *UBR5*-amplification and -duplication in the METABRIC dataset across different breast cancer subtypes. The number of tumors within each group is indicated on the top of each bar. AMP = amplification with total copy number ≥ 6; DUP = duplication with total copy number ≥ 3; Hetero_DEL = heterozygous deletion; nLOH = neutral loss-of-heterozygosity.

## References

1. Schwarz, C., Johnson, A., Kõivomägi, M., Zatulovskiy, E., Kravitz, C.J., Doncic, A., and Skotheim, J.M. (2018). A Precise Cdk Activity Threshold Determines Passage through the Restriction Point. Mol Cell 69, 253–264.e5. 10.1016/j.molcel.2017.12.017.

2. Cappell, S.D., Chung, M., Jaimovich, A., Spencer, S.L., and Meyer, T. (2016). Irreversible APCCdh1 Inactivation Underlies the Point of No Return for Cell-Cycle Entry. Cell 166, 167–180. 10.1016/j.cell.2016.05.077.

3. Moser, J., Miller, I., Carter, D., and Spencer, S.L. (2018). Control of the Restriction Point by Rb and p21. Proceedings of the National Academy of Sciences 115, E8219–E8227. 10.1073/pnas.1722446115.

4. Zetterberg, A., Larsson, O., and Wiman, K.G. (1995). What is the restriction point? Curr Opin Cell Biol 7, 835–842. 10.1016/0955-0674(95)80067-0.

5. Rubin, S.M., Sage, J., and Skotheim, J.M. (2020). Integrating Old and New Paradigms of G1/S Control. Mol Cell 80, 183–192. 10.1016/j.molcel.2020.08.020.

6. Chung, M., Liu, C., Yang, H.W., Köberlin, M.S., Cappell, S.D., and Meyer, T. (2019). Transient Hysteresis in CDK4/6 Activity Underlies Passage of the Restriction Point in G1. Molecular Cell 76, 562–573.e4. 10.1016/j.molcel.2019.08.020.

7. Liu, C., Konagaya, Y., Chung, M., Daigh, L.H., Fan, Y., Yang, H.W., Terai, K., Matsuda, M., and Meyer, T. (2020). Altered G1 signaling order and commitment point in cells proliferating without CDK4/6 activity. Nat Commun 11, 5305. 10.1038/s41467-020-18966-9.

8. Cappell, S.D., Mark, K.G., Garbett, D., Pack, L.R., Rape, M., and Meyer, T. (2018). EMI1 switches from being a substrate to an inhibitor of APC/CCDH1 to start the cell cycle. Nature 558, 313–317. 10.1038/s41586-018-0199-7.

9. Rubin, S.M. (2013). Deciphering the Rb phosphorylation code. Trends Biochem Sci 38, 12–19. 10.1016/j.tibs.2012.10.007.

10. Mittnacht, S. (1998). Control of pRB phosphorylation. Current Opinion in Genetics & Development 8, 21–27. 10.1016/S0959-437X(98)80057-9.

11. Knudsen, E.S., and Wang, J.Y. (1997). Dual mechanisms for the inhibition of E2F binding to RB by cyclin-dependent kinase-mediated RB phosphorylation. Mol Cell Biol 17, 5771–5783. 10.1128/MCB.17.10.5771.

12. Brown, V.D., Phillips, R.A., and Gallie, B.L. (1999). Cumulative Effect of Phosphorylation of pRB on Regulation of E2F Activity. Mol Cell Biol 19, 3246–3256.

13. Winston, J.T., and Pledger, W.J. (1993). Growth factor regulation of cyclin D1 mRNA expression through protein synthesis-dependent and -independent mechanisms. Mol Biol Cell 4, 1133–1144.

14. Fassl, A., Geng, Y., and Sicinski, P. (2022). CDK4 and CDK6 kinases: From basic science to cancer therapy. Science 375, eabc1495. 10.1126/science.abc1495.

15. Narasimha, A.M., Kaulich, M., Shapiro, G.S., Choi, Y.J., Sicinski, P., and Dowdy, S.F. (2014). Cyclin D activates the Rb tumor suppressor by mono-phosphorylation. eLife 3, e02872. 10.7554/eLife.02872.

16. Sanidas, I., Morris, R., Fella, K.A., Rumde, P.H., Boukhali, M., Tai, E.C., Ting, D.T., Lawrence, M.S., Haas, W., and Dyson, N.J. (2019). A Code of Mono-phosphorylation Modulates the Function of RB. Mol Cell 73, 985–1000.e6. 10.1016/j.molcel.2019.01.004.

17. Barnes, D.M., and Gillett, C.E. (1998). Cyclin D1 in Breast Cancer. Breast Cancer Res Treat 52, 1–15. 10.1023/A:1006103831990.

18. Musgrove, E.A., Caldon, C.E., Barraclough, J., Stone, A., and Sutherland, R.L. (2011). Cyclin D as a therapeutic target in cancer. Nat Rev Cancer 11, 558–572. 10.1038/nrc3090.

19. Ewen, M.E., and Lamb, J. (2004). The activities of cyclin D1 that drive tumorigenesis. Trends in Molecular Medicine 10, 158–162. 10.1016/j.molmed.2004.02.005.

20. Reissmann, P.T., Koga, H., Figlin, R.A., Holmes, E.C., Slamon, D.J., and The Lung Cancer Study Group (1999). Amplification and overexpression of the cyclin D1 and epidermal growth factor receptor genes in non-small-cell lung cancer. J Cancer Res Clin Oncol 125, 580 61–70. 10.1007/s004320050243.

21. Jiang, W., Kahn, S.M., Tomita, N., Zhang, Y.-J., Lu, S.-H., and Weinstein, I.B. (1992). Amplification and Expression of the Human Cyclin D Gene in Esophageal Cancer1. Cancer Research 52, 2980–2983.

22. Jeffreys, S.A., Becker, T.M., Khan, S., Soon, P., Neubauer, H., de Souza, P., and Powter, B. (2022). Prognostic and Predictive Value of CCND1/Cyclin D1 Amplification in Breast Cancer With a Focus on Postmenopausal Patients: A Systematic Review and Meta- Analysis. Front Endocrinol (Lausanne) 13, 895729. 10.3389/fendo.2022.895729.

23. Fukuda, T., Furuya, K., Takahashi, K., Orimoto, A., Sugano, E., Tomita, H., Kashiwagi, S., Kiyono, T., and Ishii, T. (2021). Combinatorial expression of cell cycle regulators is more suitable for immortalization than oncogenic methods in dermal papilla cells. iScience 24, 101929. 10.1016/j.isci.2020.101929.

24. Ramirez, R.D., Herbert, B.-S., Vaughan, M.B., Zou, Y., Gandia, K., Morales, C.P., Wright, W.E., and Shay, J.W. (2003). Bypass of telomere-dependent replicative senescence (M1) upon overexpression of Cdk4 in normal human epithelial cells. Oncogene 22, 433–444. 10.1038/sj.onc.1206046.

25. Spencer, S.L., Cappell, S.D., Tsai, F.-C., Overton, K.W., Wang, C.L., and Meyer, T. (2013). The proliferation-quiescence decision is controlled by a bifurcation in CDK2 activity at mitotic exit. Cell 155, 369–383. 10.1016/j.cell.2013.08.062.

26. Zatulovskiy, E., Zhang, S., Berenson, D.F., Topacio, B.R., and Skotheim, J.M. (2020). Cell growth dilutes the cell cycle inhibitor Rb to trigger cell division. Science 369, 466–471.

27. Zhang, S., Zatulovskiy, E., Arand, J., Sage, J., and Skotheim, J.M. (2022). The cell cycle inhibitor RB is diluted in G1 and contributes to controlling cell size in the mouse liver. Preprint at bioRxiv, 10.1101/2022.06.08.495371.

28. Xie, S., Zhang, S., Medeiros, G.Q.G. de, Liberali, P., and Skotheim, J. (2024). The G1/S transition in mammalian stem cells in vivo is autonomously regulated by cell size. Preprint at bioRxiv, 10.1101/2024.04.09.588781.

29. Topacio, B.R., Zatulovskiy, E., Cristea, S., Xie, S., Tambo, C.S., Rubin, S.M., Sage, J., Kõivomägi, M., and Skotheim, J.M. (2019). Cyclin D-Cdk4,6 Drives Cell-Cycle Progression via the Retinoblastoma Protein’s C-Terminal Helix. Mol Cell 74, 758–770.e4. 10.1016/j.molcel.2019.03.020.

30. Sun, X., Bizhanova, A., Matheson, T.D., Yu, J., Zhu, L.J., and Kaufman, P.D. (2017). Ki-67 Contributes to Normal Cell Cycle Progression and Inactive X Heterochromatin in p21 Checkpoint-Proficient Human Cells. Molecular and Cellular Biology 37, e00569–16. 10.1128/MCB.00569-16.

31. Zhang, M., Kim, S., and Yang, H.W. (2023). Non-canonical pathway for Rb inactivation and external signaling coordinate cell-cycle entry without CDK4/6 activity. Nat Commun 14, 7847. 10.1038/s41467-023-43716-y.

32. Broceño, C., Wilkie, S., and Mittnacht, S. (2002). RB activation defect in tumor cell lines. Proceedings of the National Academy of Sciences 99, 14200–14205. 10.1073/pnas.212519499.

33. Xie, S., and Skotheim, J.M. (2020). A G1 Sizer Coordinates Growth and Division in the Mouse Epidermis. Current Biology 30, 916–924.e2. 10.1016/j.cub.2019.12.062.

34. Sakaue-Sawano, A., Kurokawa, H., Morimura, T., Hanyu, A., Hama, H., Osawa, H., Kashiwagi, S., Fukami, K., Miyata, T., Miyoshi, H., et al. (2008). Visualizing Spatiotemporal Dynamics of Multicellular Cell-Cycle Progression. Cell 132, 487–498. 10.1016/j.cell.2007.12.033.

35. Xia, C., Fan, J., Emanuel, G., Hao, J., and Zhuang, X. (2019). Spatial transcriptome profiling by MERFISH reveals subcellular RNA compartmentalization and cell cycle-dependent gene expression. Proceedings of the National Academy of Sciences 116, 19490–19499. 10.1073/pnas.1912459116.

36. Yang, H.W., Cappell, S.D., Jaimovich, A., Liu, C., Chung, M., Daigh, L.H., Pack, L.R., Fan, Y., Regot, S., Covert, M., et al. (2020). Stress-mediated exit to quiescence restricted by increasing persistence in CDK4/6 activation. eLife 9, e44571. 10.7554/eLife.44571.

37. Dang, F., Nie, L., Zhou, J., Shimizu, K., Chu, C., Wu, Z., Fassl, A., Ke, S., Wang, Y., Zhang, J., et al. (2021). Inhibition of CK1ε potentiates the therapeutic efficacy of CDK4/6 inhibitor in breast cancer. Nat Commun 12, 5386. 10.1038/s41467-021-25700-6.

38. Repetto, M.V., Winters, M.J., Bush, A., Reiter, W., Hollenstein, D.M., Ammerer, G., Pryciak, P.M., and Colman-Lerner, A. (2018). CDK and MAPK Synergistically Regulate Signaling Dynamics via a Shared Multi-site Phosphorylation Region on the Scaffold Protein Ste5. Molecular Cell 69, 938–952.e6. 10.1016/j.molcel.2018.02.018.

39. Burke, J.R., Liban, T.J., Restrepo, T., Lee, H.-W., and Rubin, S.M. (2014). Multiple Mechanisms for E2F Binding Inhibition by Phosphorylation of the Retinoblastoma Protein C- Terminal Domain. Journal of Molecular Biology 426, 245–255. 10.1016/j.jmb.2013.09.031.

40. Deshaies, R.J., Emberley, E.D., and Saha, A. (2010). Control of cullin-ring ubiquitin ligase activity by nedd8. Subcell Biochem 54, 41–56. 10.1007/978-1-4419-6676-6_4.

41. Luo, Z., Pan, Y., Jeong, L.S., Liu, J., and Jia, L. (2012). Inactivation of the Cullin (CUL)- RING E3 ligase by the NEDD8-activating enzyme inhibitor MLN4924 triggers protective autophagy in cancer cells. Autophagy 8, 1677–1679. 10.4161/auto.21484.

42. Sdek, P., Ying, H., Zheng, H., Margulis, A., Tang, X., Tian, K., and Xiao, Z.-X.J. (2004). The central acidic domain of MDM2 is critical in inhibition of retinoblastoma-mediated suppression of E2F and cell growth. J Biol Chem 279, 53317–53322. 10.1074/jbc.M406062200.

43. Sdek, P., Ying, H., Chang, D.L.F., Qiu, W., Zheng, H., Touitou, R., Allday, M.J., and Xiao, Z.-X.J. (2005). MDM2 Promotes Proteasome-Dependent Ubiquitin-Independent Degradation of Retinoblastoma Protein. Molecular Cell 20, 699–708. 10.1016/j.molcel.2005.10.017.

44. Tomita, T., Huibregtse, J.M., and Matouschek, A. (2020). A masked initiation region in retinoblastoma protein regulates its proteasomal degradation. Nat Commun 11, 2019. 10.1038/s41467-020-16003-3.

45. Wang, Y., Zheng, Z., Zhang, J., Wang, Y., Kong, R., Liu, J., Zhang, Y., Deng, H., Du, X., and Ke, Y. (2015). A Novel Retinoblastoma Protein (RB) E3 Ubiquitin Ligase (NRBE3) Promotes RB Degradation and Is Transcriptionally Regulated by E2F1 Transcription Factor *. Journal of Biological Chemistry 290, 28200–28213. 10.1074/jbc.M115.655597.

46. Boyer, S.N., Wazer, D.E., and Band, V. (1996). E7 Protein of Human Papilloma Virus-16 Induces Degradation of Retinoblastoma Protein through the Ubiquitin-Proteasome Pathway1. Cancer Research 56, 4620–4624.

47. Uchida, C., Miwa, S., Kitagawa, K., Hattori, T., Isobe, T., Otani, S., Oda, T., Sugimura, H., Kamijo, T., Ookawa, K., et al. (2005). Enhanced Mdm2 activity inhibits pRB function via ubiquitin-dependent degradation. EMBO J 24, 160–169. 10.1038/sj.emboj.7600486.

48. Liu, H., Wang, J., Liu, Y., Hu, L., Zhang, C., Xing, B., and Du, X. (2018). Human U3 protein14a is a novel type ubiquitin ligase that binds RB and promotes RB degradation depending on a leucine-rich region. Biochimica et Biophysica Acta (BBA) - Molecular Cell Research 1865, 1611–1620. 10.1016/j.bbamcr.2018.08.016.

49. Sun, E., and Zhang, P. (2021). RNF12 Promotes Glioblastoma Malignant Proliferation via Destructing RB1 and Regulating MAPK Pathway. J Healthc Eng 2021, 4711232. 10.1155/2021/4711232.

50. de Vivo, A., Sanchez, A., Yegres, J., Kim, J., Emly, S., and Kee, Y. (2019). The OTUD5– UBR5 complex regulates FACT-mediated transcription at damaged chromatin. Nucleic Acids Research 47, 729–746. 10.1093/nar/gky1219.

51. Gudjonsson, T., Altmeyer, M., Savic, V., Toledo, L., Dinant, C., Grøfte, M., Bartkova, J., Poulsen, M., Oka, Y., Bekker-Jensen, S., et al. (2012). TRIP12 and UBR5 Suppress Spreading of Chromatin Ubiquitylation at Damaged Chromosomes. Cell 150, 697–709. 10.1016/j.cell.2012.06.039.

52. Tsai, J.M., Aguirre, J.D., Li, Y.-D., Brown, J., Focht, V., Kater, L., Kempf, G., Sandoval, B., Schmitt, S., Rutter, J.C., et al. (2023). UBR5 forms ligand-dependent complexes on chromatin to regulate nuclear hormone receptor stability. Molecular Cell 0. 10.1016/j.molcel.2023.06.028.

53. Mark, K.G., Kolla, S., Aguirre, J.D., Garshott, D.M., Schmitt, S., Haakonsen, D.L., Xu, C., Kater, L., Kempf, G., Martínez-González, B., et al. (2023). Orphan quality control shapes network dynamics and gene expression. Cell 0. 10.1016/j.cell.2023.06.015.

54. Azuma, H., Paulk, N., Ranade, A., Dorrell, C., Al-Dhalimy, M., Ellis, E., Strom, S., Kay, M.A., Finegold, M., and Grompe, M. (2007). Robust expansion of human hepatocytes in Fah-/-/Rag2-/-/Il2rg-/- mice. Nat Biotechnol 25, 903–910. 10.1038/nbt1326.

55. Grompe, M. (2017). Fah Knockout Animals as Models for Therapeutic Liver Repopulation. Adv Exp Med Biol 959, 215–230. 10.1007/978-3-319-55780-9_20.

56. Zhu, M., Lu, T., Jia, Y., Luo, X., Gopal, P., Li, L., Odewole, M., Renteria, V., Singal, A.G., Jang, Y., et al. (2019). Somatic Mutations Increase Hepatic Clonal Fitness and Regeneration in Chronic Liver Disease. Cell 177, 608–621.e12. 10.1016/j.cell.2019.03.026.

57. Curtis, C., Shah, S.P., Chin, S.-F., Turashvili, G., Rueda, O.M., Dunning, M.J., Speed, D., Lynch, A.G., Samarajiwa, S., Yuan, Y., et al. (2012). The genomic and transcriptomic architecture of 2,000 breast tumours reveals novel subgroups. Nature 486, 346–352. 10.1038/nature10983.

58. Rueda, O.M., Sammut, S.-J., Seoane, J.A., Chin, S.-F., Caswell-Jin, J.L., Callari, M., Batra, R., Pereira, B., Bruna, A., Ali, H.R., et al. (2019). Dynamics of breast-cancer relapse reveal late-recurring ER-positive genomic subgroups. Nature 567, 399–404. 10.1038/s41586-019-1007-8.

59. Hehl, L.A., Horn-Ghetko, D., Prabu, J.R., Vollrath, R., Vu, D.T., Berrocal, D.A.P., Mulder, M.P.C., Noort, G.J. van der H. van, and Schulman, B.A. (2023). Structural snapshots along K48-linked ubiquitin chain formation by the HECT E3 UBR5. Preprint at bioRxiv, 10.1101/2023.06.06.543850.

60. Hodáková, Z., Grishkovskaya, I., Brunner, H.L., Bolhuis, D.L., Belačić, K., Schleiffer, A., Kotisch, H., Brown, N.G., and Haselbach, D. (2022). Cryo-EM structure of the chain- elongating E3 ligase UBR5. Preprint at bioRxiv, 10.1101/2022.11.03.515015.

61. Wang, F., He, Q., Zhan, W., Yu, Z., Finkin-Groner, E., Ma, X., Lin, G., and Li, H. (2023). Structure of the human UBR5 E3 ubiquitin ligase. Structure 31, 541–552.e4. 10.1016/j.str.2023.03.010.

62. Jiang, W., Wang, S., Xiao, M., Lin, Y., Zhou, L., Lei, Q., Xiong, Y., Guan, K.-L., and Zhao, S. (2011). Acetylation regulates gluconeogenesis by promoting PEPCK1 degradation via recruiting the UBR5 ubiquitin ligase. Mol Cell 43, 33–44. 10.1016/j.molcel.2011.04.028.

63. Sanidas, I., Lee, H., Rumde, P.H., Boulay, G., Morris, R., Golczer, G., Stanzione, M., Hajizadeh, S., Zhong, J., Ryan, M.B., et al. (2022). Chromatin-bound RB targets promoters, enhancers, and CTCF-bound loci and is redistributed by cell-cycle progression. Mol Cell 82, 3333–3349.e9. 10.1016/j.molcel.2022.07.014.

64. Suski, J.M., Braun, M., Strmiska, V., and Sicinski, P. (2021). Targeting cell-cycle machinery in cancer. Cancer Cell 39, 759–778. 10.1016/j.ccell.2021.03.010.

65. Finn, R.S., Martin, M., Rugo, H.S., Jones, S., Im, S.-A., Gelmon, K., Harbeck, N., Lipatov, O.N., Walshe, J.M., Moulder, S., et al. (2016). Palbociclib and Letrozole in Advanced Breast Cancer. New England Journal of Medicine 375, 1925–1936. 10.1056/NEJMoa1607303.

66. Hortobagyi, G.N., Stemmer, S.M., Burris, H.A., Yap, Y.-S., Sonke, G.S., Paluch-Shimon, S., Campone, M., Blackwell, K.L., André, F., Winer, E.P., et al. (2016). Ribociclib as First-Line Therapy for HR-Positive, Advanced Breast Cancer. N Engl J Med 375, 1738–1748. 10.1056/NEJMoa1609709.

67. Sherr, C.J., Beach, D., and Shapiro, G.I. (2016). Targeting CDK4 and CDK6: From Discovery to Therapy. Cancer Discovery 6, 353–367. 10.1158/2159-8290.CD-15-0894.

68. Garrido-Castro, A.C., and Goel, S. (2017). CDK4/6 Inhibition in Breast Cancer: Mechanisms of Response and Treatment Failure. Curr Breast Cancer Rep 9, 26–33. 10.1007/s12609-017-0232-0.

69. Spring, L.M., Wander, S.A., Andre, F., Moy, B., Turner, N.C., and Bardia, A. (2020). Cyclin- dependent kinase 4 and 6 inhibitors for hormone receptor-positive breast cancer: past, present, and future. The Lancet 395, 817–827. 10.1016/S0140-6736(20)30165-3.

70. Malumbres, M., Sotillo, R., Santamaría, D., Galán, J., Cerezo, A., Ortega, S., Dubus, P., and Barbacid, M. (2004). Mammalian cells cycle without the D-type cyclin-dependent kinases Cdk4 and Cdk6. Cell 118, 493–504. 10.1016/j.cell.2004.08.002.

71. Kozar, K., Ciemerych, M.A., Rebel, V.I., Shigematsu, H., Zagozdzon, A., Sicinska, E., Geng, Y., Yu, Q., Bhattacharya, S., Bronson, R.T., et al. (2004). Mouse development and cell proliferation in the absence of D-cyclins. Cell 118, 477–491. 10.1016/j.cell.2004.07.025.

72. Regev, A., Teichmann, S.A., Lander, E.S., Amit, I., Benoist, C., Birney, E., Bodenmiller, B., Campbell, P., Carninci, P., Clatworthy, M., et al. (2017). The Human Cell Atlas. eLife 6, e27041. 10.7554/eLife.27041.

73. Abraham, K.J., Khosraviani, N., Chan, J.N.Y., Gorthi, A., Samman, A., Zhao, D.Y., Wang, M., Bokros, M., Vidya, E., Ostrowski, L.A., et al. (2020). Nucleolar RNA polymerase II drives ribosome biogenesis. Nature 585, 298–302. 10.1038/s41586-020-2497-0.

74. Charni-Natan, M., and Goldstein, I. (2020). Protocol for Primary Mouse Hepatocyte Isolation. STAR Protoc 1, 100086. 10.1016/j.xpro.2020.100086.

75. Jin, Y., Anbarchian, T., Wu, P., Sarkar, A., Fish, M., Peng, W.C., and Nusse, R. (2022). Wnt signaling regulates hepatocyte cell division by a transcriptional repressor cascade. Proceedings of the National Academy of Sciences 119, e2203849119. 10.1073/pnas.2203849119.

76. Berenson, D.F., Zatulovskiy, E., Xie, S., and Skotheim, J.M. (2019). Constitutive expression of a fluorescent protein reports the size of live human cells. MBoC 30, 2985–2995. 10.1091/mbc.E19-03-0171.

77. Lucchesi, C., Mohibi, S., and Chen, X. (2021). Measuring Translation Efficiency by RNA Immunoprecipitation of Translation Initiation Factors. Methods Mol Biol 2267, 73–79. 10.1007/978-1-0716-1217-0_5.

78. Liu, X., Yan, J., and Kirschner, M.W. (2022). Beyond G1/S regulation: how cell size homeostasis is tightly controlled throughout the cell cycle? Preprint at bioRxiv, 10.1101/2022.02.03.478996.

79. Pereira, B., Chin, S.-F., Rueda, O.M., Vollan, H.-K.M., Provenzano, E., Bardwell, H.A., Pugh, M., Jones, L., Russell, R., Sammut, S.-J., et al. (2016). The somatic mutation profiles of 2,433 breast cancers refine their genomic and transcriptomic landscapes. Nat Commun 7, 11479. 10.1038/ncomms11479.

